# Loop-Mediated Isothermal Amplification Assays for the Detection of Antimicrobial Resistance Elements in *Vibrio cholerae*

**DOI:** 10.1101/2024.06.29.601342

**Authors:** Daniel Antonio Negrón, Shipra Trivedi, Nicholas Tolli, David Ashford, Gabrielle Melton, Stephanie Guertin, Katharine Jennings, Bryan D. Necciai, Shanmuga Sozhamannan, Bradley W. Abramson

## Abstract

The bacterium *Vibrio cholerae* causes diarrheal illness and can acquire genetic material leading to multiple drug resistance (MDR). Rapid detection of resistance-conferring mobile genetic elements helps avoid the prescription of ineffective antibiotics. Colorimetric loop-mediated isothermal amplification (LAMP) assays provide a rapid and cost-effective means for detection at point-of-care, but it can be difficult to design primer sets, determine target specificity, and interpret subjective color changes. We developed an algorithm for the *in silico* design and evaluation of LAMP assays within the open-source PCR Signature Erosion Tool (PSET) and a computer vision application for the quantitative analysis of colorimetric outputs. As an example, we generated new LAMP assays targeting drug resistance in *V. cholerae* and evaluated existing ones based on *in silico* target specificity and *in vitro* testing. Improvements in the design and testing of LAMP assays, with heightened target specificity and a simple analysis platform, increase utility for in-field applications.

## 1 Introduction

Cholera, the diarrheal illness caused by ingestion of *Vibrio cholerae*, affects 1.3 to 4 million people worldwide, where 1 in 10 develop severe symptoms caused by dehydration according to the US Center for Disease Control and Prevention (CDC) (1). In 2022, 80 countries reported data on cholera to the World Health Organization (WHO). Of these, 44 countries reported 472,697 cases and 2,349 deaths, a case-fatality rate (CFR) of 0.5% in both outbreaks and as imported cases (2). Prevention and treatment options are available, such as the oral cholera vaccines (OCVs) and intravenous rehydration therapy, which, together with good hygiene practices, are optimal for reducing spread, morbidity, and mortality (3,4). However, due to vaccine shortages, antibiotics may still be recommended to help reduce the severity and duration of symptoms, especially in severe or special cases, such as pregnant individuals or those with comorbidities. Unfortunately, widespread antibiotic use has led to increasing reports of antimicrobial resistance (AMR) and multidrug resistance (MDR) in *V. cholerae* (5–7). To slow the acquisition of further resistance among cholera strains, prophylactic antibiotic treatment is not advised, and targeting the prescribed antibiotic as precisely as possible is ideal.

Frontline medical providers can optimize medical treatments based on point-of-care diagnostics, which can reduce lengthy analysis times and help improve patient outcomes. For example, molecular diagnostics such as polymerase chain reaction (PCR) and quantitative PCR (qPCR) assays quickly and reliably amplify target genetic material to detect the presence of targeted pathogenic organisms, but field usage is limited since they require bulky or expensive thermocycler instruments. Recently, loop-mediated isothermal amplification (LAMP) assays have become popular for bacterial and viral sample analyses; they are quicker than PCR, do not require expensive equipment, and provide a visual colorimetric readout of positive detection (8–12).

However, the use of LAMP assays faces challenges that slow adoption and limit their usefulness. First, primer panels for LAMP assays are more difficult to design than PCR and qPCR assays, since the primer set solution space is constrained by additional thermodynamic, length, and distance requirements. Several software applications, like the NEB® LAMP Primer Design Tool or Primer Explorer, can help design primers based on user-input target sequences, but these applications do not provide specificity metrics for the resulting assay with respect to taxonomy (13–15). Second, once a successful assay panel has been designed, the resulting colorimetric change must be assessed visually for the presence or absence of the pathogen. Subjective human interpretation of color can be unreliable, time-consuming, and limited in sensitivity (8,16). To address this need, recently, visible colorimetric measurements using smartphone or tablet software and cameras (i.e., smartphones) have been developed (17–19).

The PCR Signature Erosion Tool (PSET) provides an *in silico* method for determining primer binding sites relative to taxonomically related sequences and detects mutations in primers used for PCR or qPCR assays (20,21). For this study, we created LAMP assays for *V. cholerae* detection, beginning with improvements to PSET, and culminating with a robust, quantitative determination of the assay outcome using computer vision. The new LAMP workflow module for PSET can generate assays or predict performance with respect to large reference sequence databases, such as the NCBI nucleotide (nt) database. Running the tool periodically can identify potential assay failures as newly evolved sequences are deposited into databases such as GenBank or GISAID (20). Additionally, we developed a LAMP assay design tool based on the open source Primer3 program as an alternative to web-based programs, leveraging multicore architecture. We then performed end-to-end design of novel LAMP assays together with *in silico* evaluation of existing ones. Finally, we validated the LAMP assay designs *in vitro* and developed a method to computationally determine the colorimetric changes of the LAMP assay using a smartphone camera. All code is publicly available at https://github.com/biolaboro/PSET.

## 2 Methods

### 2.1 Design *in silico*

A LAMP assay design script was written in Python (v3.11.4) with bindings to the Primer3 PCR design tool (v2.6.1) via the primer3-py (v2.6.0) API (22–25). A Snakemake workflow coordinates execution of the script, which first runs Primer3 to identify primer pairs and optional loop candidates (26). It uses the Primer3 configuration file system to guide the nested search for inner primers and loops with additional parameters for inter-primer distances, strandedness, and thermodynamic constraints. A procedure then evaluates every primer pair and loop primer combination. Accordingly, there is an initial search for the outer F3/B3 primers and inner F2/B2 primers. The next four searches calculate the F1c, B1c, LF, and LB candidates separately. Nonoverlapping combinations of F2-LF-F1c and B1c-LB-B2 are then evaluated based on F1c-B1c distance constraints and for 5’/3’ stability in terms of ΔG. The final LAMP 5’/3’ assay definition indicates primers and loops with square brackets and parentheses on the amplicon. Additionally, each primer, loop, FIP, and BIP component is individually defined in a resulting JSON object with an associated assay penalty score equal to the sum of Primer3 penalties.

### 2.2 Evaluation *in silico*

An alignment-based workflow implements the LAMP assay evaluation within the PSET framework. The initial phase queries the amplicon against a BLAST+ database (27), collecting a list of accessions with sequence identity and coverage above a parameterized threshold. The next phase realigns each component primer and loop separately, using the glsearch36 tool of the FASTA suite (28) for a global-local alignment to guarantee full query coverage. A filtering step removes results with alignment below a parameterized identity threshold. The last phase checks for nested primer and loop order, arrangement, and inter-primer/loop distances. A confusion matrix is output with respect to each subject accession, where calls are based on subject taxonomy and whether primer and loop sequences arranged correctly while exceeding identity threshold. The taxonomy check determines whether the NCBI Taxonomy identifier of the subject is equal to or a descendant of any of the identifiers in the set of assay targets. Accordingly, the taxonomy and arrangement-alignment evaluation yield the true/false and positive/negative component of the confusion matrix call.

### 2.3 *In vitro* LAMP assay testing

All LAMP primers (Table S1) were ordered from IDT and reconstituted at 100μM in sterile water. FIP and BIP primers were diluted to 16μM, F3 and B3 primers were diluted to 2μM, and LF and LB primers were diluted to 4μM to make a 10X primer master mix. Positive control gene Block (gBlock) fragments were ordered from IDT and reconstituted at 10ng/μL. Serial (1:10) dilutions were performed in molecular grade water to achieve the concentrations of 10pg/μL through 1fg/μL that were tested. LAMP reagents were ordered from NEB (WarmStart Colorimetric LAMP 2X Master Mix cat# M1800S). The LAMP reaction mix was prepared using 12.5μL of WarmStart Colorimetric LAMP 2X Master Mix, 2.5μL of the reaction specific LAMP 10X primer mix, and 9μL of molecular grade water per well of a 96-well reaction plate (Applied Biosystems cat# N8010560). The final primer concentrations were 1.6μM for FIP and BIP, 0.2μM for F3 and B3, and 0.4μM for LF and LB. 1μL of the desired dilution of each gene fragment was then added to the corresponding well and 1μL of molecular grade water was added to no template control wells. The plate was incubated at 65°C for 30 minutes in a thermocycler (Applied Biosystems TFS-2720) following LAMP 2X Master Mix manufacture guidelines. After the incubation period, the plate was allowed to cool to room temperature. Final LAMP assay images were captured with a smartphone (Samsung Galaxy S10e) with default camera settings.

### 2.4 LAMPvision image analysis

The LAMPvision prototype python script uses the NetworkX (3.0), NumPy (v1.26.4), OpenCV (v4.9.0), Pandas (2.2.0), Scikit-Learn (1.5.0) packages to analyze the LAMP assay images (19,29–33). First, the algorithm filters the image to restrict colors within the HSV (hue, saturation, value) color range between (0, 85, 100) and (179, 255, 255), which correspond to pinkish reds and yellows of the LAMP assay. Next, the Hough circle transform procedure detects circles within the grayscale representation of the image. This step returns the location and size of each circle detected and is meant to locate plate wells. Overlapping circles are combined into a bigger one based on radius and the new location is set to the centroid. Next, the circles are binned into a grid index based on horizontal and vertical overlap. Accordingly, a 1-D clustering step uses the mean-shift algorithm to calculate the minimum cluster value based on the pairwise distances of the circles. This requires that the well plate have at least two non-empty adjacent wells. The resulting value reflects the average distance between adjacent wells. This is done so that the output corresponds to each well plate index, even if a row or column is empty. Finally, the mean RGB color of each circle is output with corresponding well index. An R script (v4.4.0) calculated the color difference between each reaction and the corresponding no-template negative control and rendered plots with the ggplot2 library (3.5.1) and related extensions (34–38). These color differences across replicates (n = 4 for each assay sample) were then plotted with the relative copies per well. Uniquely, the graph plots each point as the extracted well pixels from the original image. Raw images, metadata, and scripts are available in the analysis.zip archive file.

## 3 Results

### 3.1 LAMP assay primer design and PSET analysis of *V. cholerae* AMR genes

A functional LAMP assay requires careful primer design, with considerations such as amplicon length, primer spacing, primer orientation, and primer annealing temperatures, while also requiring specificity to the target sequence (10). Here we designed and incorporated a LAMP primer design tool into PSET for rapid and automated determination of primer erosion that occurs when mutations in the template strand mismatch with the primer cause failure of a previously working assay (Figure 1). Briefly, the LAMP assay design algorithm uses Primer3 to determine the F3/B3 primer pair, followed by the internal primer pairs F2/F1c and B1c/B2 which are concatenated into FIP and BIP, respectively. Within the latter pairs, additional optional primers are chosen to become Loop-F (LF) and Loop-B (LB) primers, which can help speed-up nucleic acid amplification during the assay. The final LAMP assay is designed based on appropriate spacing of these primer sets as well as the annealing temperature differences to create a constraint-based LAMP assay for the target sequence.

**Figure 1.**
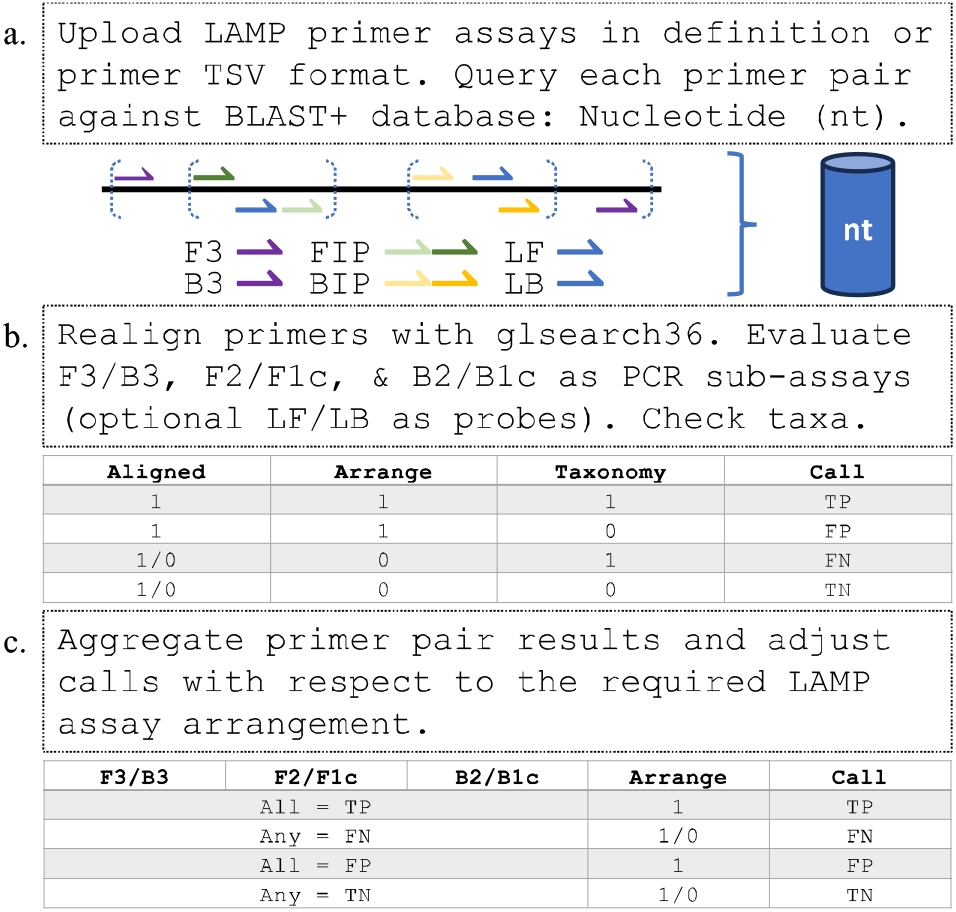
Conceptual design of LAMP primers. a) The user loads assays as LAMP primers or in an assay definition format for b) blasting to find target sequences and validate primer spacing. Appropriate primer alignments, primer arrangements, and taxonomy targets are calculated. c) A confusion matrix is generated for each assay to describe the target specificity.

PSET evaluates LAMP assays by analyzing primer alignment and arrangement against BLAST database taxa to determine *in silico* target specificity. The previous version of PSET determined PCR or qPCR assays which have 2 or 3 primer sets respectively. Therefore, an algorithm was required for LAMP assays containing 6-8 total primers which are not in a typical PCR primer orientation (Figure 1). Since the FIP/BIP primers create a loop structure through an inverted hairpin, their sequences are a combination of disjointed forward and reverse primers F2/F1c or B2/B1c respectively. An algorithm was developed to reduce the FIP and BIP primers to their counterparts. Briefly, a BLAST search queries the F3/B3 primers to retrieve a template sequence for the FIP/BIP primers, which are then aligned to it. The hits are then evaluated where the FIP is expected to have at least one alignment in the forward direction of 18-22nt and at least one alignment of 18-22nt in the reverse direction roughly 10-50nts up or downstream.

For integration into the PSET evaluation routine, LAMP assays are treated as three pseudo-PCR/qPCR assays representing amplicons from the pairs of primer for F3/B3, F2/F1c with LF denoted as a probe, and B2/B1c with LB denoted as a probe. Each one is evaluated for correct orientation and spacing. The final output is a cumulative “call” of the LAMP assays that reflects primer alignment, arrangement, and subject sequence taxonomy. True Positive (TP) LAMP assay targets must have all pseudo-PCR/qPCR assays return individual TP calls to the same subject. If any pseudo-PCR/qPCR assays return a False Negative (FN) then the LAMP assay is considered an FN, even if primers are oriented correctly, which demonstrates that the overall LAMP assay would not effectively target the subject sequence of interest.

We demonstrated PSET’s taxonomy detection feature using previously published PCR, qPCR, or LAMP primer sets targeting *V. cholerae* AMR genes, while additionally designing novel LAMP primer sets (Table S1). These primer sets were compared to NCBI’s nt database targeting NCBI Taxonomy identifier 666 (*V. cholerae*) using PSET (Table S2). In general, previously published assays largely targeted *V. cholerae*, however the nt database does not provide a means of distinguishing submissions with AMR. Therefore, some of the AMR targets like *gyrA, tcp*, and *ompU* genes result in high FP hits, suggesting that greater than 80% of the submitted nucleotide sequences do not contain these specific AMR genes (Table S2). Therefore, we created a custom taxonomy database using *V. cholerae* sequences from Microbial Browser for Identification of Genetic and Genomic Elements (MicroBIGG-E) that are associated with AMR classes and genes. Given that PSET operates using NCBI’s taxonomy database, we created a system to import new taxonomies based on AMR classes from MicroBIGG-E (Table S3) and tested the LAMP assays designed here (Table S4). 34 of 43 assays showed an AMR class specificity of greater than 97%. However, the assays with low specificity targeted tetracycline and quinolone where these AMR classes have multiple genes for conferring resistance. This is likely due to gene-specific targeting since several are capable of conferencing resistance to the same class.

### 3.2 In vitro testing with independent quantitative analysis methods

Primers were designed using the design principles described above and the target gene fragments were synthesized based on deposited sequences from MicroBIGG-E (22). The samples were serially diluted, producing a 10-fold dilution curve for each assay alongside a sample that did not contain any template as a negative control (Figure 2). Four independent replicates were incubated for 30min at 60°C. These LAMP colorimetric assays produce a pH dependent color shift in response to nucleic acid amplification where negative samples are red and positive samples are yellow. A quantitative visual interpretation of the color change was determined by two individuals by eye (Table S5) as well as an automated interpretation based on image capture and hexadecimal-valued color distance (Table S6 and S7). Nine of the 16 assays produced positive signals in the highest concentration template reaction while the no-template control was still negative, and four assays remained negative at all DNA template concentrations suggesting no amplification occurred. Three of the assays had amplification in all reactions, including the negative control, and it is unclear at this time why this has occurred.

**Figure 2.**
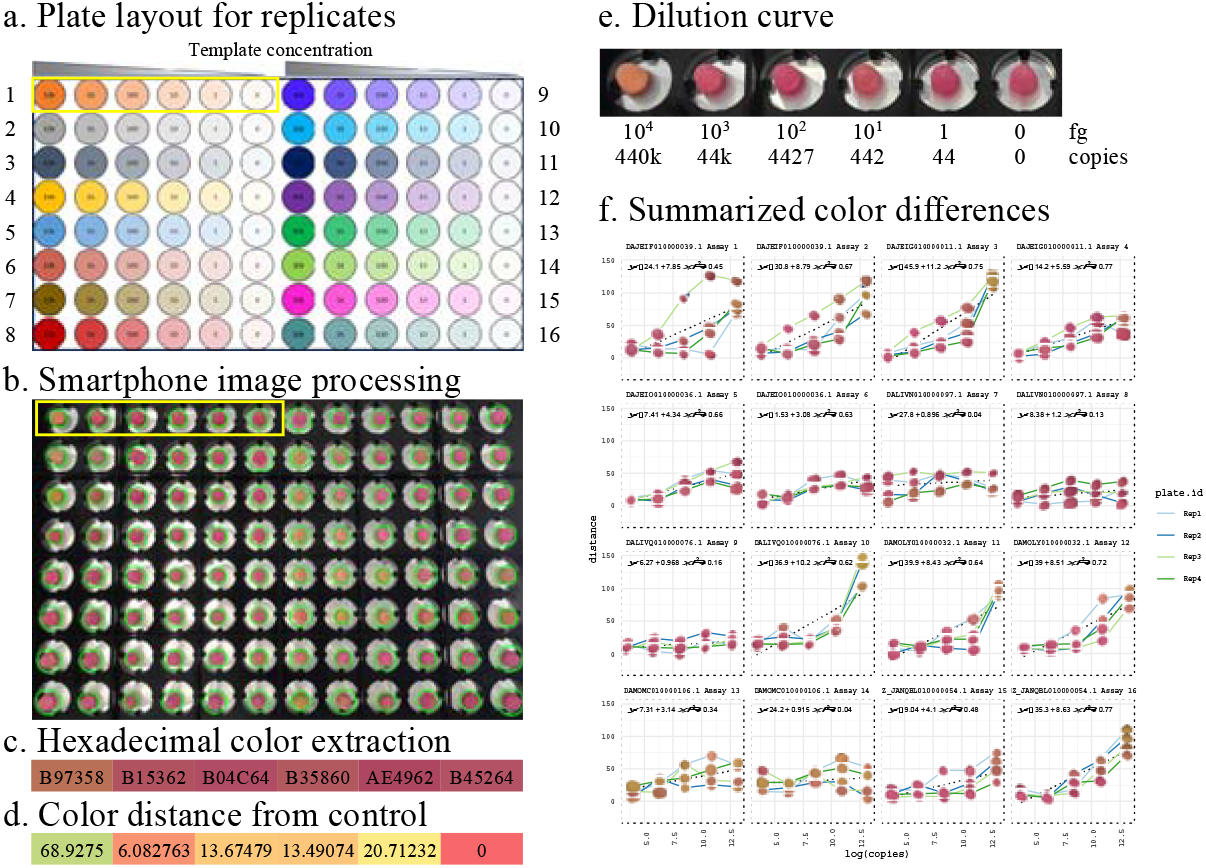
in vitro LAMP assay analysis. a) Plate layout scheme for the replicates (n=4) performed. Each assay consisted of 5 samples with template and a no template control. A total of 16 unique assays were run on one plate and performed four separate days. b) Image of the completed LAMP assay with detected wells within the green circles. c) Extracted hexadecimal color values. d) Each sample’s color difference to the no template control was calculated. e) One assay displayed zoomed in describing dilutions and copies of the template used. f) The color difference was plotted relative to the copies of the template for each reaction for the four replicates. The color dots on the plots are the extracted well color and shape.

Visual analysis of LAMP assays showed detection of 5 AMR gene targets for *aph(6), varG, floR, qnrVC5*, and *almG*, but analysis by visual interpretation is subjective, time consuming, and difficult to interpret individual wells on a plate. Therefore, automated image analysis was employed. Using a cellphone to capture images of the plates, we determined color differences of the sample to their cognizant no-template control (Figure 2d). These color differences across replicates (n=4 for each reaction) are then plotted with the relative copies per well (Figure 2f). Each point on the graph consists of the extracted well pixels from the original image.

The slope was calculated from the LAMPvision color distance plots to determine if assays have a positive signal trajectory with increasing template concentration and it was found that a slope threshold of 4.0 may be used to determine if positive signal is produced, but further testing is required (Figure 3a). Next, we determined the correlation of the methods for the LAMP assay visual analysis and the LAMPvision tool (Figure 3). Eight assays showed a high degree of correlation (>0.8 Pearson correlation) between LAMP vision and the visual quantitative analysis (Figure 3a). Correlations were not calculated for four assays because the visual analysis was not able to determine any positive signal and all values were zero by visual analysis (Table S5). Three assays did not show a color change but had a strong correlation (0.5-0.8 Pearson correlation) and one assay had a negative correlation. These results suggest there is a strong agreement in the visual analysis and LAMPvision analysis when there is positive signal of the assay.

**Figure 3.**
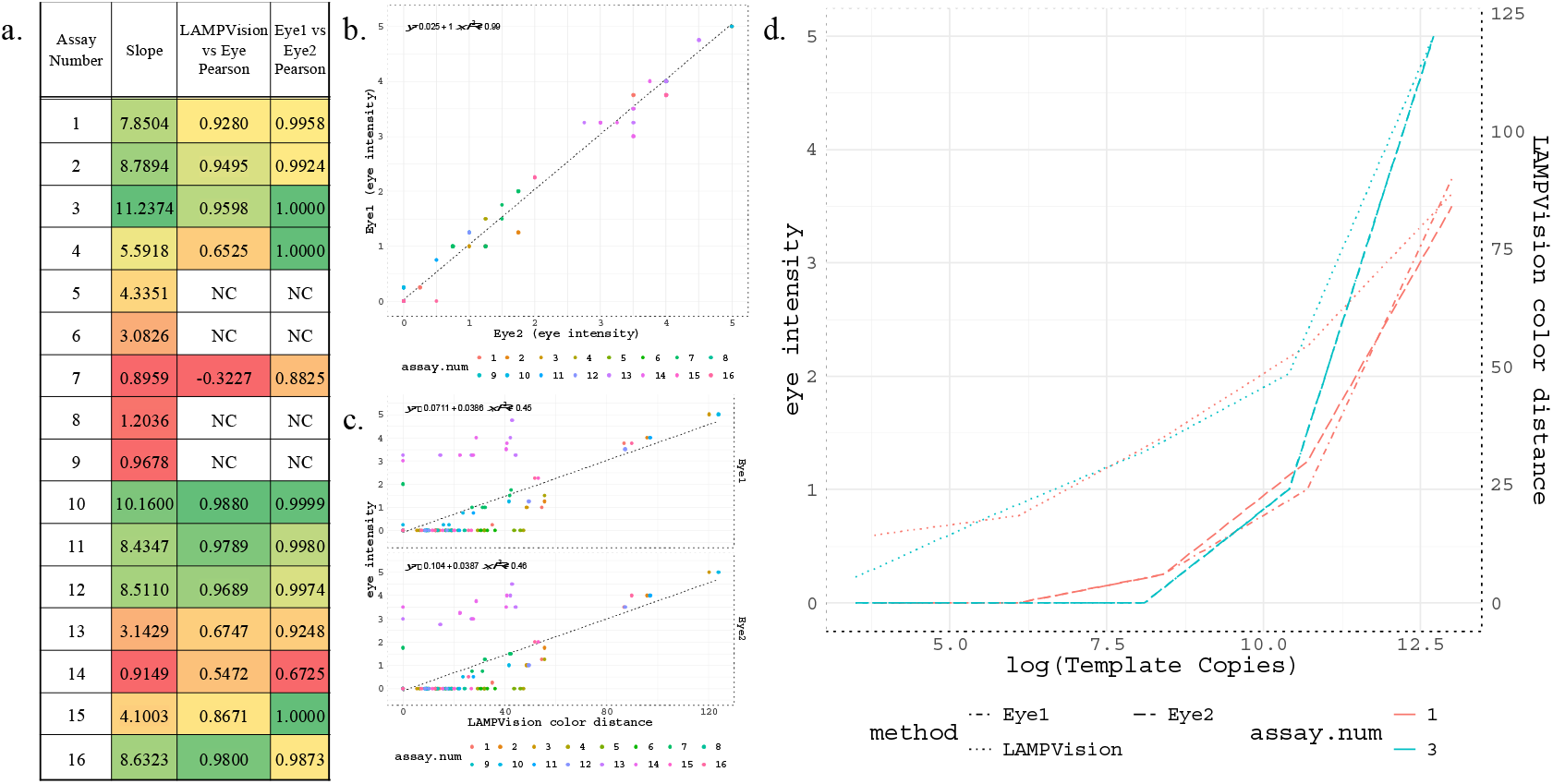
Comparison of visual interpretation to quantitative computer LAMPvision methods. a) The Pearson correlation was calculated for the human visual determination of color shift on a scale of 0-5 and compared to the LAMPvision methods described here. NC indicates non-computable due to standard deviation of 0. b) A comparison describing the subjective nature of two human individuals’ interpretation of the color change. c) Another comparison of color scores by visual analysis (Eye1, top; Eye2, bottom) versus computer vision analysis. d) A plot showing the difference in color change detected visually and by computer vision methods. Computer vision methods enhance the limit of detection.

## 4 Discussion

Here we show the *in vitro* application of *in silico* LAMP assay design incorporated into the Primer Signature Erosion Tool (PSET) and a computer vision algorithm to interpret complex plate-based colorimetric LAMP assays. By combining all these aspects, we can rapidly design, test, and validate novel LAMP assays targeting AMR genes while also determining suitability of previously published LAMP assays for target specificity. The rapid design and validation of these LAMP assays allows individuals to develop vast arrays of assays targeting numerous genes, signature sequences of species, or variants of concern. Here we validated the code to produce novel LAMP assays and detect AMR synthetic gene sequences. Although this work primarily focused on *in silico* aspects, limited *in vitro* data show great promise for on-demand assay design and testing in far-forward scenarios, applications, and use-cases.

Colorimetric LAMP assays as a diagnostic have been approved by the FDA for some target strains, such as *Salmonella* (39) or SARS-CoV-2 (40), via Emergency Use Authorization (EUA) and demonstrates their utility. The reliability of a nucleic acid detection method that is easy to use and interpret will help track novel and recurring pathogens and can aid in the surveillance of non-infectious environmental DNA (eDNA). Using the software designed here, new LAMP assays can be designed, analyzed, and output into an ordering form for rapid prototyping as demonstrated here (Figure 2).

*In silico* target specificity was calculated for 66 previously published LAMP, PCR, and qPCR assays from 27 publications targeting AMR genes in *V. cholerae* (Table S1). Additionally, we designed 43 new LAMP assays with the described LAMP primer design and PSET integration tools. These assays were tested via PSET for their *in silico* target specificity against NCBI’s nt database (Table S2) and the *V. cholerae* subset of the MicroBIGG-E database (Table S3).

LAMP colorimetric assays are used as rapid, point-of-care diagnostics. However, due to the non-specific pH-based color changes, they cannot be multiplexed in a single well. An array of individual colorimetric LAMP assays can be used to detect multiple targets, but then the interpretation of numerous assays becomes problematic. Here we designed a useful computer vision tool, LAMPvision, to detect wells and quantify results reliably across several replicates (n=4). Other studies have used wavelength shifts in absorption spectra or smartphone applications, but those studies required expensive machines or complicated setups to correct for lighting (39, 41). LAMPvision uses an internal negative control placed next to the samples to serve as a color normalization factor. However, there were observable differences in saturation across the images that may introduce signal noise. These may be mitigated by performing additional background correction processes. Further testing is required to account for dynamic lighting or photo angle.

Several of the LAMP assays were not functional and required revisions to the ordering form of the code, initially. For instance, the B3 primer was ordered in the wrong orientation relative to the assay due to a bug in the output script. However, the assays were still functional and produced positive signals. Interestingly, the utility of the B3 primer is to facilitate removal of the FIP/BIP primers during amplification. This could be one reason these assays have high limits of detection in terms of copy number. Further refinement of these metrics will presumably result in a higher rate of LAMP assay success. Similarly, since only synthetic DNA controls were used in these experiments, further characterization on live samples is required. Furthermore, much more stringent controls and validation steps are required for clinical usage.

Color image analysis is difficult to compare to human eye perception and additional testing may be required to validate the color distance calculations. In this method, we used red, green, and blue color values which do not account for color intensity and perhaps a color difference calculation using hue, saturation, lightness (value; HSL, HSV) is better. Since RGB hexadecimal values can fluctuate with brightness, intensity correction methods may be required for validation and could be implemented as negative control wells strategically interspersed. Likewise, we did not determine a specific negative control color value and instead used the associated assays’ no-template control. Accordingly, having one negative control does not provide a means to calculate negative standard deviations for thresholding positive signals. In future applications, all sample negative controls may be utilized.

Overall, we present a means of rapidly prototyping new LAMP assays with target specificity including *in vivo* tests with computer vision analysis. Their point-of-care usage can help combat increasing antimicrobial resistance as LAMP assays become easier to access and interpret for individuals. As these tests become robust and well validated, they will have greater utility in field testing to control and mitigate cholera outbreaks. As outlined in the Global Roadmap 2030.2, intensified strategic and systematic use of rapid diagnostic tests followed by laboratory confirmation will contribute to reaching the targets of the Ending Cholera: (WER, 2023, 98, 431-452)

## Supporting information

Analysis data and scripts...

## Acknowledgment

Funding for this work was provided by JPEO (CBRND), JPL-EB, DBPAO and laboratory work was funded by the Noblis Sponsored Research program for internal R&D.

## Author Contributions

Conceptualization: DN, DA, SS, and BA. Methodology: DN, ST, SS, and BA. Software: DN, ST, and BA. Validation: DN, NT, GM, and BA. Formal analysis: DN, and BA. Investigation: DN, NT, GM, and BA. Resources: KJ, SS, and BA. Data Curation: DN, and BA. Writing - Original Draft: DN, NT, and BA. Writing - Review & Editing: DN, NT, DA, SG, KJ, SS, and BA. Visualization: DN, and BA. Supervision: KJ, SS, and BA. Project administration: DN, DA, KJ, BN, SS, and BA. Funding acquisition: KJ, BN, SS, and BA.

## Disclaimer

The views expressed in this manuscript are those of the authors and do not necessarily reflect the official policy or position of the JPEO-CBRND, the Department of Defense, or the U.S. Government. This work was prepared as part of the author(s) official duties. Title 17 U.S.C. § 105 provides that ‘Copyright protection under this title is not available for any work of the United States Government.’ Title 17 U.S.C. §101 defines U.S. Government work as work prepared by a military service member or employee of the U.S. Government as part of that person’s official duties. References to non-federal entities or their products do not constitute or imply Department of Defense or Army endorsement of any company, product or organization. Funding for this study was provided and executed by the Joint Program Executive Office for Chemical, Biological, Radiological and Nuclear Defense’s (JPEO-CBRND) Joint Project Lead for CBRND Enabling Biotechnologies (JPL CBRND EB) on behalf of the Department of Defense’s Chemical and Biological Defense Program.

## References

1. Ali M, Nelson AR, Lopez AL, Sack DA. Updated Global Burden of Cholera in Endemic Countries. PLOS Neglected Tropical Diseases. 2015 Jun 4;9(6):e0003832.

2. WHO. Cholera, 2022. WER. 2023 Sep 22;98(38):431–43.

3. WHO. Cholera vaccines: WHO position paper – August 2017. WER. 2017 Aug 25;92(34):477–500.

4. CDC. Cholera. 2024 [cited 2024 May 21]. Treating Cholera. Available from: https://www.cdc.gov/cholera/treatment/index.html

5. Das B, Verma J, Kumar P, Ghosh A, Ramamurthy T. Antibiotic resistance in Vibrio cholerae: Understanding the ecology of resistance genes and mechanisms. Vaccine. 2020 Feb 29;38:A83–92.

6. Yuan X hui, Li Y mei, Vaziri AZ, Kaviar VH, Jin Y, Jin Y, et al. Global status of antimicrobial resistance among environmental isolates of Vibrio cholerae O1/O139: a systematic review and meta-analysis. Antimicrob Resist Infect Control. 2022 Dec;11(1):62.

7. Verma J, Bag S, Saha B, Kumar P, Ghosh TS, Dayal M, et al. Genomic plasticity associated with antimicrobial resistance in Vibrio cholerae. Proc Natl Acad Sci USA. 2019 Mar 26;116(13):6226–31.

8. Kashir J, Yaqinuddin A. Loop mediated isothermal amplification (LAMP) assays as a rapid diagnostic for COVID-19. Med Hypotheses. 2020 Aug;141:109786.

9. Zhao X, Zeng Y, Yan B, Liu Y, Qian Y, Zhu A, et al. A novel extraction-free dual HiFi-LAMP assay for detection of methicillin-sensitive and methicillin-resistant Staphylococcus aureus. Microbiology Spectrum. 2024 Feb 20;0(0):e04133–23.

10. Notomi T, Mori Y, Tomita N, Kanda H. Loop-mediated isothermal amplification (LAMP): principle, features, and future prospects. J Microbiol. 2015 Jan 1;53(1):1–5.

11. Bao H, Zhao Y, Wang Y, Xu X, Shi J, Zeng X, et al. Development of a Reverse Transcription Loop-Mediated Isothermal Amplification Method for the Rapid Detection of Subtype H7N9 Avian Influenza Virus. BioMed Research International. 2014 Feb 6;2014:e525064.

12. Leroy AG, Persyn E, Gibaud SA, Crémet L, Le Turnier P, Benhamida M, et al. Assessment of a Multiplex LAMP Assay (Eazyplex® CSF Direct M) for Rapid Molecular Diagnosis of Bacterial Meningitis: Accuracy and Pitfalls. Microorganisms. 2021 Sep 1;9(9):1859.

13. New England Biolabs. NEB® LAMP Primer Design Tool [Internet]. [cited 2024 May 24]. Available from: https://lamp.neb.com/#!/

14. Eiken Chemical Co., Ltd. LAMP primer designing software PrimerExplorer [Internet]. [cited 2024 May 24]. Available from: https://primerexplorer.jp/e/index.html

15. Akhmetzianova LU, Davletkulov TM, Sakhabutdinova AR, Chemeris AV, Gubaydullin IM, Garafutdinov RR. LAMPrimers iQ: New primer design software for loop-mediated isothermal amplification (LAMP). Analytical Biochemistry. 2024 Jan 1;684:115376.

16. Ahn SJ, Baek YH, Lloren KKS, Choi WS, Jeong JH, Antigua KJC, et al. Rapid and simple colorimetric detection of multiple influenza viruses infecting humans using a reverse transcriptional loop-mediated isothermal amplification (RT-LAMP) diagnostic platform. BMC Infectious Diseases. 2019 Aug 1;19(1):676.

17. Colbert AJ, Lee DH, Clayton KN, Wereley ST, Linnes JC, Kinzer-Ursem TL. PD-LAMP smartphone detection of SARS-CoV-2 on chip. Analytica Chimica Acta. 2022 Apr 22;1203:339702.

18. García-Bernalt Diego J, Fernández-Soto P, Márquez-Sánchez S, Santos Santos D, Febrer-Sendra B, Crego-Vicente B, et al. SMART-LAMP: A Smartphone-Operated Handheld Device for Real-Time Colorimetric Point-of-Care Diagnosis of Infectious Diseases via Loop-Mediated Isothermal Amplification. Biosensors. 2022 Jun;12(6):424.

19. Heithoff DM, Barnes L, Mahan SP, Fox GN, Arn KE, Ettinger SJ, et al. Assessment of a Smartphone-Based Loop-Mediated Isothermal Amplification Assay for Detection of SARS-CoV-2 and Influenza Viruses. JAMA Netw Open. 2022 Jan 4;5(1):e2145669.

20. Negrón DA, Kang J, Mitchell S, Holland MY, Wist S, Voss J, et al. Impact of SARS-CoV-2 Mutations on PCR Assay Sequence Alignment. Frontiers in Public Health [Internet]. 2022 [cited 2024 Feb 23];10. Available from: https://www.frontiersin.org/journals/public-health/articles/10.3389/fpubh.2022.889973

21. Sozhamannan S, Holland MY, Hall AT, Negrón DA, Ivancich M, Koehler JW, et al. Evaluation of Signature Erosion in Ebola Virus Due to Genomic Drift and Its Impact on the Performance of Diagnostic Assays. Viruses. 2015 Jun;7(6):3130–54.

22. Untergasser A, Cutcutache I, Koressaar T, Ye J, Faircloth BC, Remm M, et al. Primer3—new capabilities and interfaces. Nucleic Acids Res. 2012 Aug;40(15):e115.

23. Koressaar T, Remm M. Enhancements and modifications of primer design program Primer3. Bioinformatics. 2007 May 15;23(10):1289–91.

24. Kõressaar T, Lepamets M, Kaplinski L, Raime K, Andreson R, Remm M. Primer3_masker: integrating masking of template sequence with primer design software. Bioinformatics. 2018 Jun 1;34(11):1937–8.

25. Van Rossum G, Drake FL. Python 3 reference manual. Scotts Valley, CA: CreateSpace; 2009.

26. Mölder F, Jablonski KP, Letcher B, Hall MB, Tomkins-Tinch CH, Sochat V, et al. Sustainable data analysis with Snakemake. F1000Res. 2021 Jan 18;10:33.

27. Camacho C, Coulouris G, Avagyan V, Ma N, Papadopoulos J, Bealer K, et al. BLAST+: architecture and applications. BMC Bioinformatics. 2009;10(1):421.

28. Pearson WR, Lipman DJ. Improved tools for biological sequence comparison. Proc Natl Acad Sci U S A. 1988 Apr;85(8):2444–8.

29. Hagberg A, Swart PJ, Schult DA. Exploring network structure, dynamics, and function using NetworkX [Internet]. Los Alamos National Laboratory (LANL), Los Alamos, NM (United States); 2008 Jan [cited 2024 May 22]. Report No.: LA-UR-08-05495; LA-UR-08-5495. Available from: https://www.osti.gov/biblio/960616

30. Harris CR, Millman KJ, Van Der Walt SJ, Gommers R, Virtanen P, Cournapeau D, et al. Array programming with NumPy. Nature. 2020 Sep 17;585(7825):357–62.

31. Pandas development. pandas-dev/pandas: Pandas [Internet]. Zenodo; 2020. Available from: 10.5281/zenodo.3509134

32. McKinney W. Data Structures for Statistical Computing in Python. In: Walt S van der, Millman J, editors. Proceedings of the 9th Python in Science Conference. 2010. p. 56–61.

33. Pedregosa F, Varoquaux G, Gramfort A, Michel V, Thirion B, Grisel O, et al. Scikit-learn: Machine Learning in Python. Journal of Machine Learning Research. 2011;12(85):2825–30.

34. R Core Team. R: A language and environment for statistical computing [Internet]. Vienna, Austria; 2020. Available from: https://www.R-project.org/

35. Wickham H. ggplot2: Elegant graphics for data analysis [Internet]. Springer-Verlag New York; 2016. Available from: https://ggplot2.tidyverse.org

36. Yu G. ggimage: Use Image in “ggplot2” [Internet]. 2023. Available from: https://CRAN.R-project.org/package=ggimage

37. Aphalo PJ. ggpmisc: Miscellaneous Extensions to “ggplot2” [Internet]. 2024. Available from: https://CRAN.R-project.org/package=ggpmisc

38. Campitelli E. ggnewscale: Multiple Fill and Colour Scales in “ggplot2” [Internet]. 2024. Available from: https://CRAN.R-project.org/package=ggnewscale

39. Confirmation of Salmonella Isolates by Loop-Mediated Isothermal Amplification (LAMP).

40. Color SARS-CoV-2 RT-LAMP Diagnostic Assay - EUA Summary. 2021;

41. Papadakis G, Pantazis AK, Fikas N, Chatziioannidou S, Tsiakalou V, Michaelidou K, et al. Portable real-time colorimetric LAMP-device for rapid quantitative detection of nucleic acids in crude samples. Sci Rep. 2022 Mar 8;12(1):3775.

