## Supplementary material for "Loop-Mediated Isothermal Amplification Assays for the Detection of Antimicrobial Resistance Elements in *Vibrio cholerae*": Analysis data and scripts...: figures.R

````
# ---
# title: "Figures"
# author: "Daniel Antonio Negrón"
# output: html_document
# ---
  
# ```{r packages}
library(bitops)
library(dplyr)
library(ggplot2)
library(ggimage)
library(ggpmisc)
library(ggnewscale)
library(tidyr)
library(tibble)
# ```

# ```{r globals}
tmp <- tempdir()

ls.prefix <- (
  setNames(
    sapply(c(1, 2, 3, 6, 6, 9, 12, 15, 18, 21, 24, 27, 30), \(ele) 10^-ele),
    paste0(c("d", "c", "m", "μ", "u", "n", "p", "f", "a", "z", "y", "r", "q"), "g")
  )
)

to_rgb_int <- function(x) {
  x <- ifelse(substr(x, 1, 1) == "#", substr(x, 2, 7), substr(x, 1, 6))
  sapply(c(1, 3, 5), \(e) strtoi(paste0("0x", substr(x, e, e + 1))))
}

color_distance <- function(p, q) {
  # https://www.compuphase.com/cmetric.htm
  rmean <- (p[1] + q[1]) / 2
  r <- p[1] - q[1]
  g <- p[2] - q[2]
  b <- p[3] - q[3]
  sqrt((((512 + rmean) * r * r) %>>% 8) + 4 * g * g + (((767 - rmean) * b * b) %>>% 8))
}
# ```

# ```{r load}
df.mask <- (
  readxl::read_xlsx("metadata.xlsx", sheet = "mask", col_names = F, col_types = "text") %>%
    suppressMessages() %>%
    setNames(c("id", (1:(ncol(.) - 1)))) %>%
    group_by(id) %>%
    group_modify(\(.x, ...) rownames_to_column(.x, "row")) %>%
    ungroup() %>%
    pivot_longer(-c(id, row), names_to = "col") %>%
    mutate(
      contig.id = sapply(strsplit(value, " ", fixed = T), "[", 1),
      across(c(col, row), as.integer)
    ) %>%
    separate(value, c("assay.id", "concentration"), sep = " - ")
)

df.meta <- readxl::read_xlsx("metadata.xlsx", sheet = "meta", col_types = "text")

df.img <- (
  lapply(df.meta$plate.id, function(ele) {
    untar(paste0(ele, ".tar.gz"), exdir = file.path(tmp, ele))
    list.files(file.path(tmp, ele), full.names = T)
  }) %>%
    unlist() %>%
    as_tibble() %>%
    mutate(
      plate.id = basename(dirname(value)),
      name = basename(value)
    ) %>%
    separate(name, c("col", "row"), sep = "[_\\.]", remove = T, convert = T, extra = "drop") %>%
    rename(path = value)
)

df.copy <- (
  readxl::read_xlsx("metadata.xlsx", sheet = "copy", col_names = T, col_types = "text") %>%
    pivot_longer(-c(1, 2), names_to = "contig.id", values_to = "copies") %>%
    mutate(copies = as.numeric(copies))
)

df.data <- (
  lapply(df.meta$plate.id, function(ele) {
    readxl::read_xlsx(paste0(ele, ".xlsx"), col_names = F, col_types = "text") %>%
      suppressMessages() %>%
      setNames(1:ncol(.)) %>%
      rownames_to_column("row") %>%
      mutate(plate.id = ele) %>%
      pivot_longer(-c(plate.id, row), names_to = "col") %>%
      mutate(across(c(row, col), \(ele) as.integer(ele))) %>%
      left_join(df.meta, by = join_by(plate.id)) %>%
      left_join(df.mask, by = join_by(mask.id == id, col, row))
  }) %>%
    bind_rows() %>%
    left_join(
      select(filter(., concentration == "NC"), plate.id, assay.id, value),
      by = join_by(plate.id, assay.id),
      suffix = c("", ".control")
    ) %>%
    mutate(
      prefix = sub("^[0-9\\.,-]+", "", concentration),
      base = ls.prefix[prefix],
      dilution = as.numeric(gsub("[^0-9]", "", concentration)) * base,
      distance = mapply(\(p, q) color_distance(to_rgb_int(p), to_rgb_int(q)), p = value, q = value.control),
      color = paste0("#", value),
      assay.id = factor(
        assay.id,
        levels = unique(assay.id)[order(as.integer(sapply(strsplit(unique(assay.id), " "), `[`, 3)))]
      )
    ) %>%
    left_join(df.copy, by = join_by(concentration, contig.id)) %>%
    left_join(df.img, by = join_by(plate.id, col, row)) %>%
    arrange(dilution)
)
# ```

# ```{r Fig2f, fig.width=6.5}
df.data %>%
  filter(complete.cases(.)) %>%
  ggplot(aes(x = log(copies), y = distance)) +
  stat_poly_eq(use_label(c("eq", "R2")), size = 2) +
  stat_poly_line(color = "black", linetype = "dotted", linewidth = 0.5) +
  geom_path(aes(group = plate.id, color = plate.id)) +
  geom_image(aes(image = path), size = 0.1) +
  scale_color_brewer(palette = "Paired") +
  facet_wrap(~assay.id) +
  theme_minimal() +
  theme(
    axis.text.x = element_text(angle = 90),
    strip.text.y = element_text(hjust = 0.5, angle = 0),
    panel.border = element_rect(fill = NA, linetype = "dotted"),
    text = element_text(family = "mono", size = 8)
  )

ggsave("Fig2f.pdf", dpi=300)
ggsave("Fig2f.png", dpi=300)
# ```

# ```{r TableS7}
df.data %>%
  identity() %>%
  select(plate.id, row, col, distance) %>%
  arrange(plate.id, row, col) %>%
  pivot_wider(names_from = col, values_from = distance) %>%
  mutate(row = LETTERS[row]) %>%
  write.table("TableS7.tsv", quote = F, sep = "\t", row.names = F)
# ```

# ```{r eye_intensity}
df.eye <- (
  readxl::read_xlsx("TableS5.xlsx") %>%
    group_by(plate.id, eye.id) %>%
    group_modify(~ mutate(.x, row = 1:nrow(.x))) %>%
    ungroup() %>%
    select(plate.id, eye.id, row, everything())
)
# ```

# ```{r Fig3b, fig.width=6.5}
df.eye %>%
  pivot_longer(-c(plate.id, eye.id, row), names_to = "col", names_pattern = "([0-9]+)$", names_transform = list(col = as.integer)) %>%
  left_join(df.meta, by = join_by(plate.id)) %>%
  left_join(df.mask, by = join_by(mask.id == id, col, row)) %>%
  group_by(assay.id, eye.id, row, col) %>%
  summarise(value = mean(value), .groups = "drop") %>%
  pivot_wider(names_from = eye.id, values_from = value) %>%
  mutate(
    assay.num = sapply(strsplit(assay.id, " "), tail, 1),
    assay.num = factor(assay.num, levels = unique(sort(as.integer(assay.num))))
  ) %>%
  rename(
    `Eye1 (eye intensity)` = Eye1,
    `Eye2 (eye intensity)` = Eye2
  ) %>%
  ggplot(aes(`Eye2 (eye intensity)`, `Eye1 (eye intensity)`)) +
  stat_poly_eq(use_label(c("eq", "R2"))) +
  stat_poly_line(color = "black", linetype = "dotted", linewidth = 0.5) +
  geom_point(aes(color = assay.num)) +
  guides(color = guide_legend(nrow = 2, byrow = T)) +
  theme_minimal() +
  theme(
    legend.position = "bottom",
    panel.border = element_rect(fill = NA, linetype = "dotted"),
    text = element_text(family = "mono", size = 16)
  )

ggsave("Fig3b.pdf", dpi = 300)
ggsave("Fig3b.png", dpi = 300)
# ```

# ```{r Fig3c, fig.width=6.5, fig.height=6.5}
df.eye %>%
  pivot_longer(-c(plate.id, eye.id, row), names_to = "col", names_pattern = "([0-9]+)$", names_transform = list(col = as.integer)) %>%
  left_join(df.meta, by = join_by(plate.id)) %>%
  left_join(df.mask, by = join_by(mask.id == id, col, row)) %>%
  group_by(assay.id, eye.id, row, col) %>%
  summarise(value = mean(value), .groups = "drop") %>%
  left_join(
    summarise(group_by(df.data, assay.id, row, col), distance = mean(distance), .groups = "drop"),
    by = join_by(assay.id, row, col)
  ) %>%
  mutate(
    assay.num = sapply(strsplit(assay.id, " "), tail, 1),
    assay.num = factor(assay.num, levels = unique(sort(as.integer(assay.num))))
  ) %>%
  rename(eye = value) %>%
  ggplot(aes(distance, eye)) +
  stat_poly_eq(use_label(c("eq", "R2"))) +
  stat_poly_line(color = "black", linetype = "dotted", linewidth = 0.5) +
  geom_point(aes(color = assay.num)) +
  facet_grid(eye.id ~ .) +
  scale_y_continuous(breaks = 0:5) +
  guides(color = guide_legend(nrow = 2, byrow = T)) +
  xlab("LAMPVision color distance") +
  ylab("eye intensity") +
  theme_minimal() +
  theme(
    legend.position = "bottom",
    panel.border = element_rect(fill = NA, linetype = "dotted"),
    text = element_text(family = "mono", size = 16)
  )

ggsave("Fig3c.pdf", dpi = 300)
ggsave("Fig3c.png", dpi = 300)
# ```

# ```{r Fig3d, fig.width=6.5, fig.height=5}
df.fig3c <- (
  df.eye %>%
    pivot_longer(-c(plate.id, eye.id, row), names_to = "col", names_pattern = "([0-9]+)$", names_transform = list(col = as.integer)) %>%
    left_join(df.meta, by = join_by(plate.id)) %>%
    left_join(df.mask, by = join_by(mask.id == id, col, row)) %>%
    left_join(df.copy, by = join_by(concentration, contig.id)) %>%
    rename(method = eye.id) %>%
    select(method, assay.id, plate.id, col, row, copies, value) %>%
    bind_rows(
      select(df.data, assay.id, plate.id, col, row, copies, distance) %>%
        rename(value = distance) %>%
        mutate(method = "LAMPVision")
    ) %>%
    group_by(assay.id, method, row, col, copies) %>%
    summarise(value = mean(value), .groups = "drop") %>%
    # pivot_wider(names_from = method, values_from = value) %>%
    mutate(
      assay.num = sapply(strsplit(assay.id, " "), tail, 1),
      assay.num = factor(assay.num, levels = unique(sort(as.integer(assay.num))))
    ) %>%
    filter(
      assay.id %in% c("DAJEIF010000039.1 Assay 1", "DAJEIG010000011.1 Assay 3"),
      !is.na(copies)
    )
)

max_lamp_value <- max(with(df.fig3c, df.fig3c[method == "LAMPVision", "value"]))

df.fig3c %>%
  mutate(
    value = if_else(method == "LAMPVision", value / max_lamp_value * 5, value)
  ) %>%
  arrange(copies) %>%
  ggplot(aes(x = log(copies), y = value, color = assay.num, linetype = method)) +
  geom_path() +
  scale_y_continuous(
    name = "eye intensity",
    sec.axis = sec_axis(trans = ~ . / 5 * max_lamp_value, name = "LAMPVision color distance")
  ) +
  scale_linetype_manual(values = c(Eye1 = "dotdash", Eye2 = "longdash", LAMPVision = "dotted")) +
  xlab("log(Template Copies)") +
  guides(
    color = guide_legend(nrow = 2, byrow = T),
    linetype = guide_legend(nrow = 2, byrow = T, title = "method")
  ) +
  theme_minimal() +
  theme(
    panel.border = element_rect(fill = NA, linetype = "dotted"),
    text = element_text(family = "mono", size = 16),
    legend.position = "bottom"
  )

ggsave("Fig3d.pdf", dpi = 300)
ggsave("Fig3d.png", dpi = 300)
# ```

# ```{r correlation}
df.corr <- (
  df.eye %>%
    pivot_longer(-c(plate.id, eye.id, row), names_to = "col", names_pattern = "([0-9]+)$", names_transform = list(col = as.integer)) %>%
    left_join(df.meta, by = join_by(plate.id)) %>%
    left_join(df.mask, by = join_by(mask.id == id, col, row)) %>%
    group_by(assay.id, eye.id, row, col) %>%
    summarise(value = mean(value), .groups = "drop") %>%
    left_join(
      summarise(group_by(df.data, assay.id, row, col), distance = mean(distance), .groups = "drop"),
      by = join_by(assay.id, row, col)
    ) %>%
    mutate(
      assay.num = sapply(strsplit(assay.id, " "), tail, 1),
      assay.num = as.integer(assay.num, levels = unique(sort(as.integer(assay.num))))
    ) %>%
    rename(eye = value) %>%
    arrange(row, col)
)

df.disteye <- (
  df.corr %>%
    group_by(assay.num, row, col, distance) %>%
    summarise(eye = mean(eye), .groups = "drop") %>%
    group_by(assay.num) %>%
    group_map(function(.x, .y) {
      with(.x, 
           data.frame(
             .y, 
             dist_v_eye_pearson = cor(eye, distance, method = "pearson")
           )
      )
    }) %>%
    bind_rows()
)

df.slope <- (
  df.data %>%
    select(assay.id, row, col, distance, copies) %>%
    mutate(assay.num = as.integer(sapply(strsplit(as.character(assay.id), " "), tail, 1))) %>%
    group_by(assay.num, copies, row, col) %>%
    summarise(distance = mean(distance), .groups = "drop") %>%
    group_by(assay.num) %>%
    arrange(assay.num, row, desc(col)) %>%
    group_map(function(.x, .y) {
      data.frame(.y, slope = lm(distance ~ log(copies), data = .x)$coefficients[2])
    }) %>%
    bind_rows()
)

df.eyeeye <- (
  df.corr %>%
    group_by(assay.num, row, col, eye.id) %>%
    summarise(eye = mean(eye), .groups = "drop") %>%
    group_by(assay.num) %>%
    group_map(function(.x, .y) {
      values <- split(.x, .x$eye.id)
      data.frame(
        .y, 
        eye1_v_eye2_pearson = cor(values[[1]]$eye, values[[2]]$eye, method = "pearson")
      )
    }) %>%
    bind_rows()
)

left_join(df.disteye, df.slope, by = join_by(assay.num)) %>%
  left_join(df.eyeeye, by = join_by(assay.num)) %>%
  select(assay.num, dist_v_eye_pearson, slope, eye1_v_eye2_pearson) %>%
  write.table("Fig3a.tsv", quote = F, sep = "\t", row.names = F)
# ```
````
