## Supplementary material for "Loop-Mediated Isothermal Amplification Assays for the Detection of Antimicrobial Resistance Elements in *Vibrio cholerae*": Analysis data and scripts...: lampvision.py

```
#!/usr/bin/env python

import sys
import tarfile
from io import BytesIO
from itertools import combinations, pairwise
from pathlib import Path

import cv2 as cv
import networkx as nx
import numpy as np
import pandas as pd
from scipy import ndimage
from sklearn.cluster import MeanShift
from sklearn.neighbors import KDTree


def quantize(x, t=0.05):
    return int(np.ceil(x) if (np.ceil(x) - x) <= t else x)


def get_circles(img):
    # Apply Hough transform
    circles = cv.HoughCircles(
        cv.cvtColor(img, cv.COLOR_BGR2GRAY),
        cv.HOUGH_GRADIENT_ALT,
        1.5,
        1,
        param1=50,
        param2=0.1,
        minRadius=25,
        maxRadius=50
    )[0]

    img_copy = img.copy()
    for x, y, r in np.uint16(np.around(circles)):
        cv.circle(img_copy, (x, y), r, BGR_WHITE, -1)
    # cv.imshow("circles", img_copy)
    # cv.waitKey(0)

    D = nx.DiGraph()
    D.add_edges_from(
        (tuple(c1), tuple(c2))
        for c1, c2 in combinations(circles, 2)
        if np.linalg.norm(c1[:2] - c2[:2]) <= c1[2] + c2[2]
    )

    seen = set()
    for ele in nx.weakly_connected_components(D):
        seen |= ele
        overlaps = np.array(list(ele))
        centroid = overlaps[:, :2].mean(axis=0)
        radius = np.linalg.norm(centroid - overlaps[0, :2]) + overlaps[:, 2].max()
        # new circle
        yield (*centroid, radius)

    # original singletons
    yield from [ele for ele in circles if tuple(ele) not in seen]


BGR_BLACK = (0, 0, 0)
BGR_WHITE = (255, 255, 255)
BGR_GREEN = (0, 255, 0)

path = Path(sys.argv[1])
img = cv.imread(str(path))
img_hsv = cv.cvtColor(img, cv.COLOR_BGR2HSV)

color_mask = cv.inRange(img_hsv, (0, 85, 100), (179, 255, 255))
img_filtered = cv.bitwise_and(img, img, mask=color_mask)

# cv.imshow("filtered", img_filtered)
# cv.waitKey(0)

circles = np.array(list(get_circles(img_filtered)))

img_copy = img.copy()
for x, y, r in np.uint16(np.around(circles)):
    cv.circle(img_copy, (x, y), r, BGR_GREEN, 8, cv.LINE_AA)
# cv.imshow("circles", img_copy)
cv.imwrite(str(path.with_suffix(f".circles{path.suffix}")), img_copy)
# cv.waitKey(0)

# tree = KDTree(circles[:, :2])
# dists, neighbors = tree.query(circles[:, :2], 2)
# T = np.mean(circles[:, :2], axis=0)
# medians = []
# for i in range(45):
#     t = np.deg2rad(i)
#     R = np.array([[np.cos(t), -np.sin(t)], [np.sin(t), np.cos(t)]])
#     A = (circles[:, :2] - T) @ R + T
#     slopes = []
#     for ele in neighbors:
#         dx, dy = A[ele[0]] - A[ele[1]]
#         dx, dy = (dx, dy) if not np.isclose(dx, 0) else (dy, dx)
#         slopes.append(dy / dx)
#     medians.append(np.median(slopes))
# t = np.argmin(np.abs(medians))

# img_filtered = ndimage.rotate(img_filtered, t)
circles = np.array(list(get_circles(img_filtered)))

circles = np.uint16(np.around(circles))

img_filtered_circled = img_filtered.copy()
for e in circles:
    cv.circle(img_filtered_circled, e[:2], e[2], (255, 255, 255), -1)

x_bins = {}
y_bins = {}
for i, e in enumerate(circles):
    x, y, r = e
    for coor, bins in ((x, x_bins), (y, y_bins)):
        q1, q2 = coor - r, coor + r
        for s1, s2 in bins:
            if s1 <= q1 <= s2 or s1 <= q2 <= s2 or q1 <= s1 <= q2 or q1 <= s2 <= q2:
                bins[(s1, s2)].append(i)
                break
        else:
            bins[(q1, q2)] = [i]

X = np.array([
    ele2 - ele1
    for idx, bins in enumerate((x_bins, y_bins))
    for ele1, ele2 in pairwise((circles[val, idx].mean() for _, val in sorted(bins.items())))
]).reshape(-1, 1)
ms = MeanShift(bandwidth=None, bin_seeding=True)
ms.fit(X)
# print(ms.cluster_centers_)
d_min = ms.cluster_centers_.min()

x_min, y_min = circles[:, 0].min(), circles[:, 1].min()
x_max, y_max = circles[:, 0].max(), circles[:, 1].max()
nrow, ncol = 1 + int((y_max - y_min) / d_min), 1 + int((x_max - x_min) / d_min)
wrow, wcol = len(str(nrow)), len(str(ncol))

df_rgb = pd.DataFrame(None, index=list(range(nrow)), columns=list(range(ncol)), dtype=str)

with tarfile.open(path.with_suffix(".tar.gz"), "w:gz") as tar:
    for i, e in enumerate(circles):
        x, y, r = e
        img_filtered_circled = img_filtered.copy()
        cv.circle(img_filtered_circled, (x, y), r, BGR_WHITE, -1)
        white_mask_2d = np.all((img_filtered_circled == BGR_WHITE), axis=-1)
        idx = img_filtered[white_mask_2d] != BGR_BLACK
        values = [ele for ele in img_filtered[white_mask_2d] if not np.array_equal(ele, BGR_BLACK)]
        mean = np.around(np.mean(values, axis=0)).astype(np.uint8)
        color = "{:02X}{:02X}{:02X}".format(mean[2], mean[1], mean[0])
        col, row = quantize((x - x_min) / d_min), quantize((y - y_min) / d_min)
        df_rgb.at[row, col] = color

        img_filtered_circled = img_filtered.copy()[y - r:y + r, x - r:x + r]
        # cv.circle(img_filtered_circled, np.array(img_filtered_circled.shape[:2][::-1]) // 2, r, BGR_GREEN, 1)
        # Make a True/False mask of pixels whose BGR values sum to more than zero
        alpha = np.sum(img_filtered_circled, axis=-1) > 0
        # Convert True/False to 0/255 and change type to "uint8" to match "na"
        alpha = np.uint8(alpha * 255)
        # Stack new alpha layer with existing image to go from BGR to BGRA, i.e. 3 channels to 4 channels
        img_filtered_circled = np.dstack((img_filtered_circled, alpha))
        fileobj = BytesIO(cv.imencode(".png", img_filtered_circled, [cv.IMWRITE_PNG_COMPRESSION, 9])[1])
        tarinfo = tarfile.TarInfo(f"{(col+1):0{wcol}}_{(row+1):0{wrow}}.png")
        tarinfo.size = fileobj.getbuffer().nbytes
        tar.addfile(tarinfo=tarinfo, fileobj=fileobj)
        # print(row, col, d_min, y, y_min, (y - y_min) / d_min, x, x_min, (x - x_min) / d_min, sep="\t")

df_rgb.style.map(lambda value: None if pd.isna(value) else f"background-color: #{value};").to_excel(path.with_suffix(".xlsx"), header=False, index=False)

# cv.destroyAllWindows()
```
