## Supplementary material for "Loop-Mediated Isothermal Amplification Assays for the Detection of Antimicrobial Resistance Elements in *Vibrio cholerae*": Analysis data and scripts...: README.txt

Install PSET from https://github.com/biolaboro/PSET and activate the conda environment. 
Then run the figures.sh script.
Note: some of the values may change slightly as the analysis may use different seeds when clustering within the lampvision.py prototype script.
