## Supplementary figures and images for "Loop-Mediated Isothermal Amplification Assays for the Detection of Antimicrobial Resistance Elements in *Vibrio cholerae*"

### Fig2f.pdf

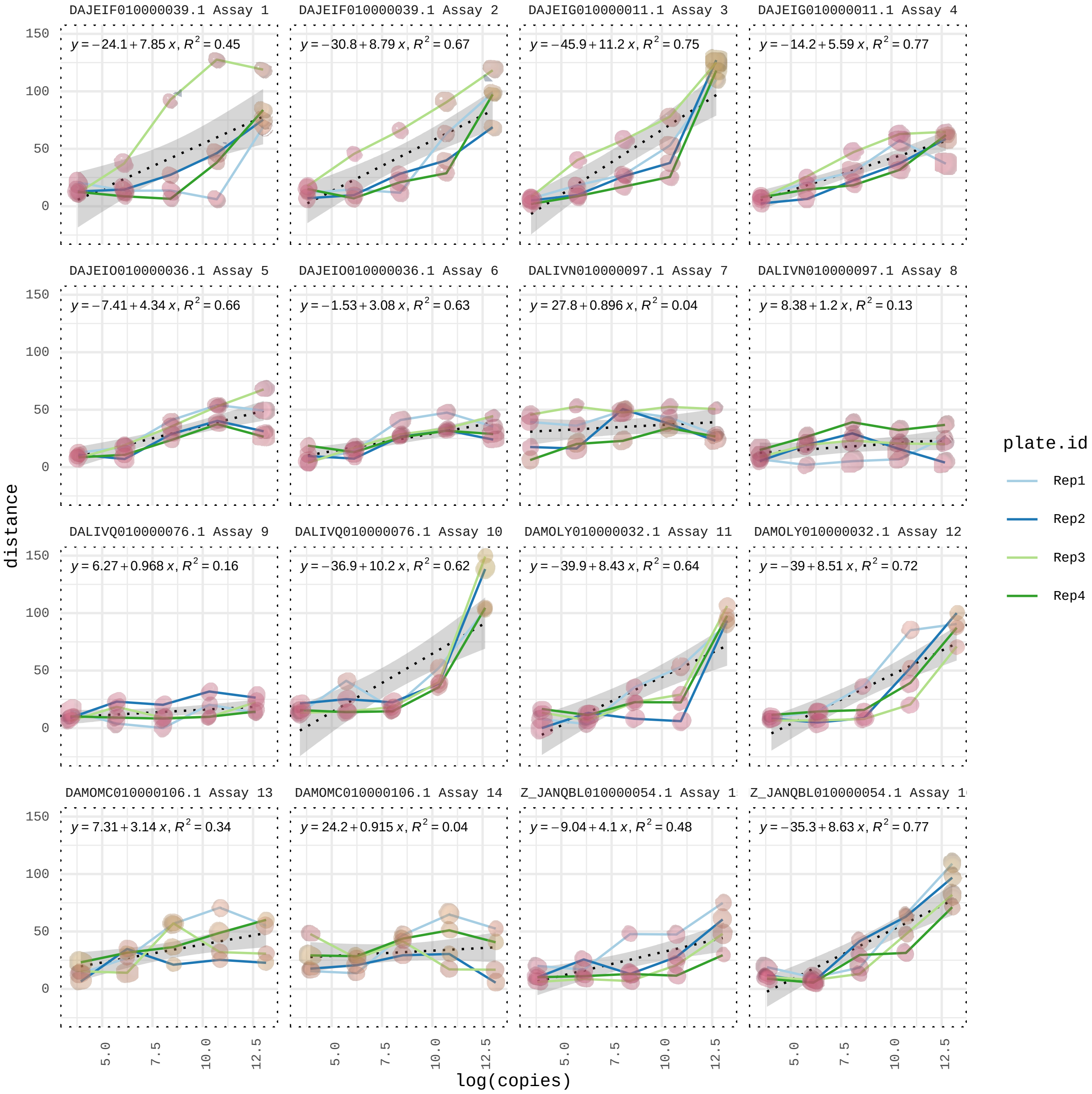

### Fig2f.png

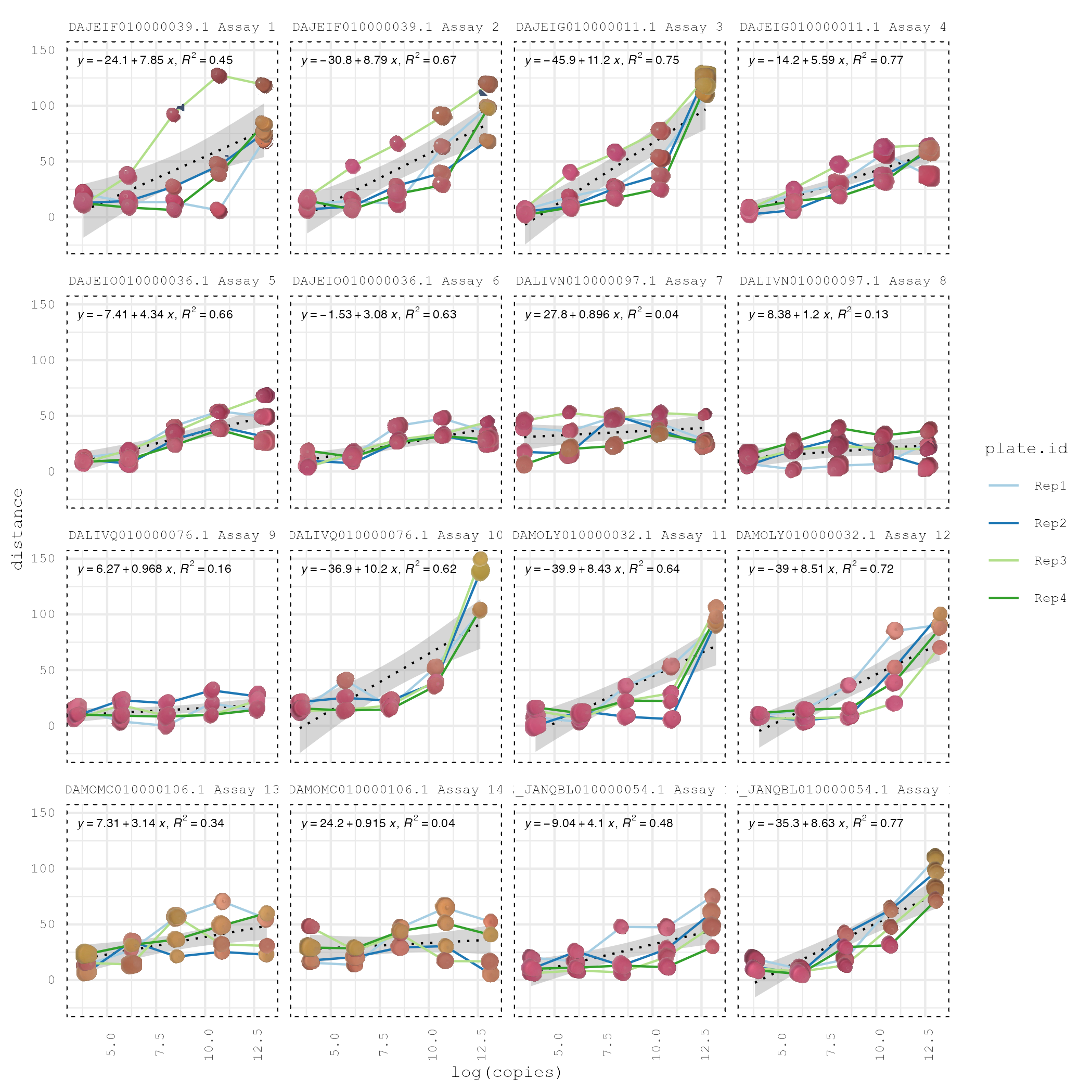

### Fig3b.pdf

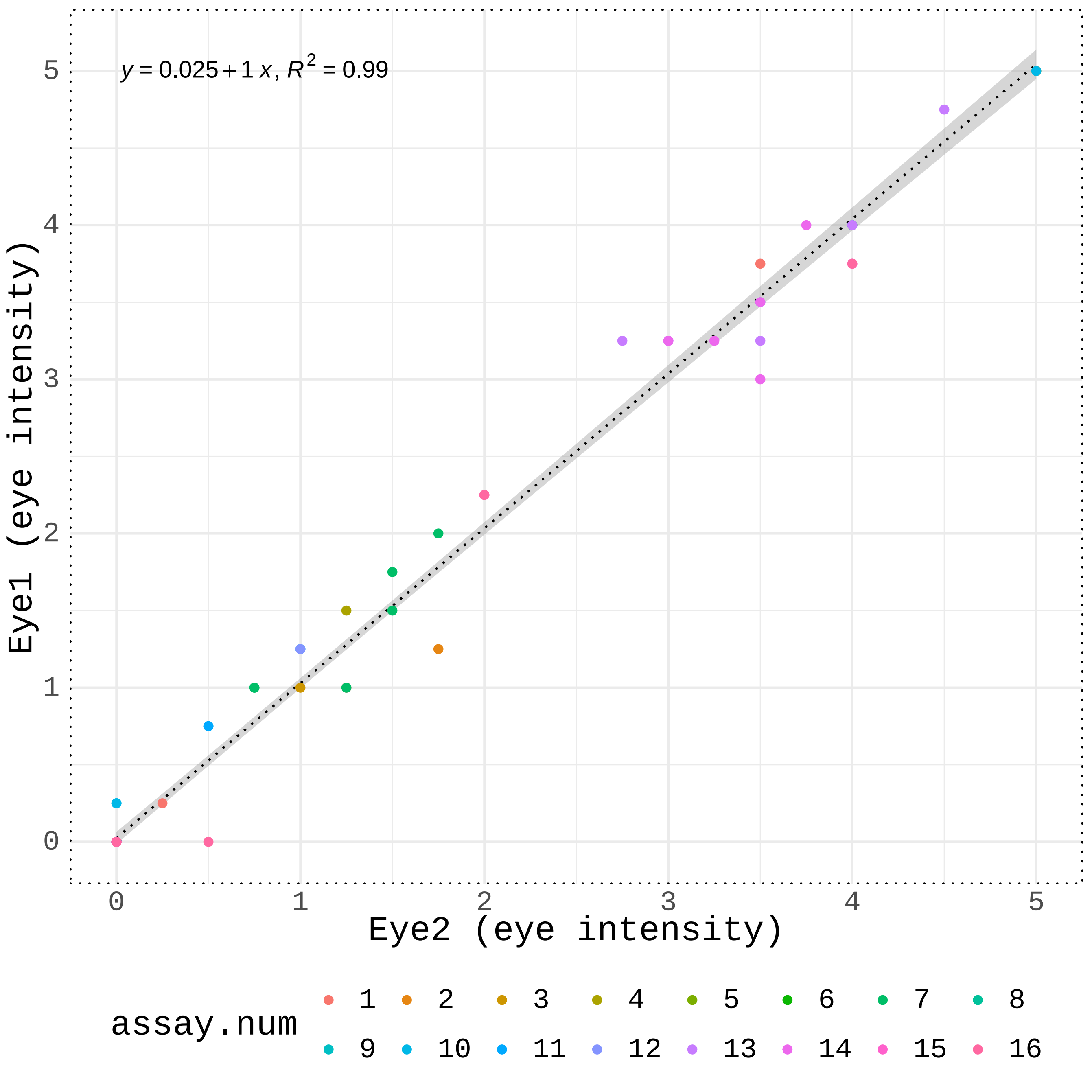

### Fig3b.png

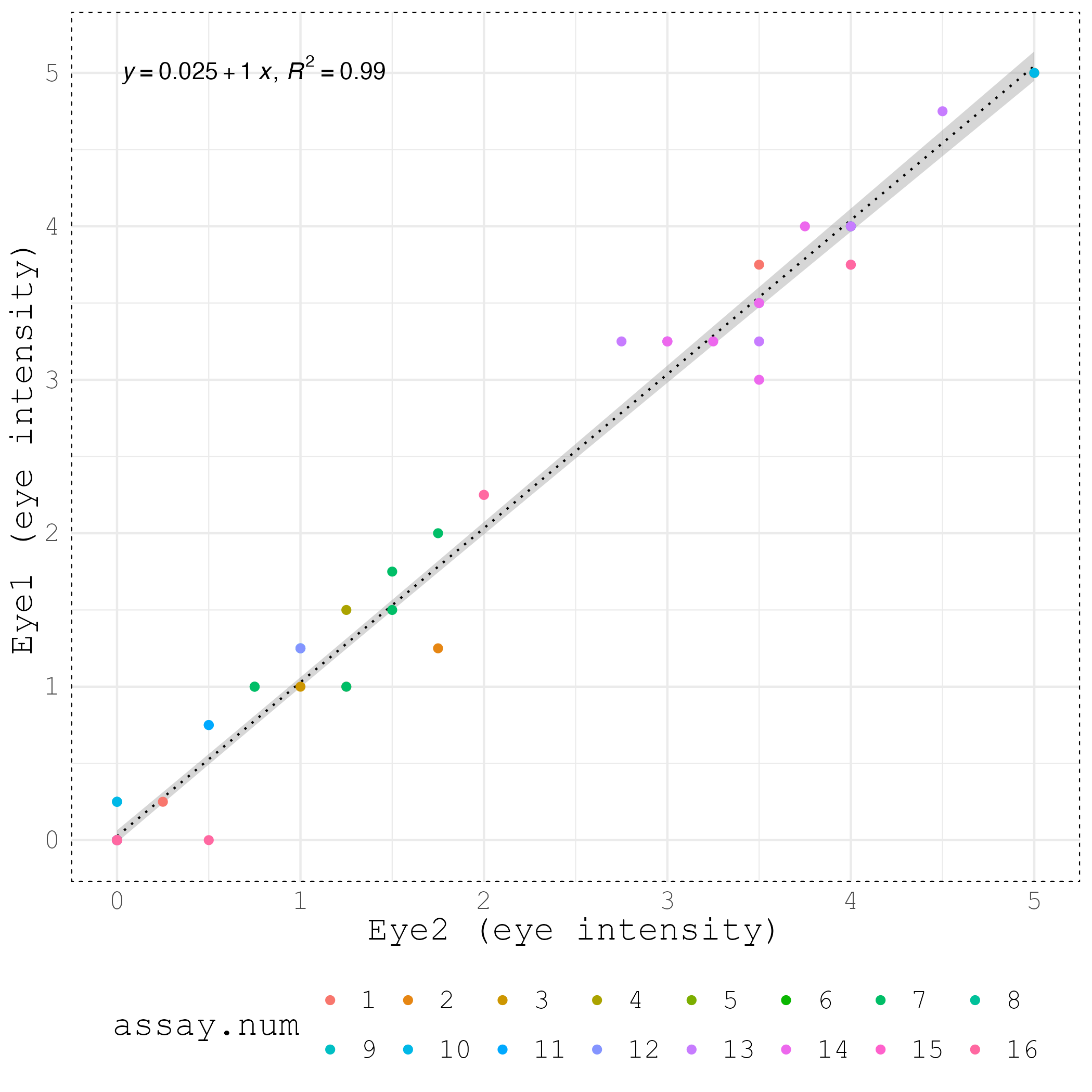

### Fig3c.pdf

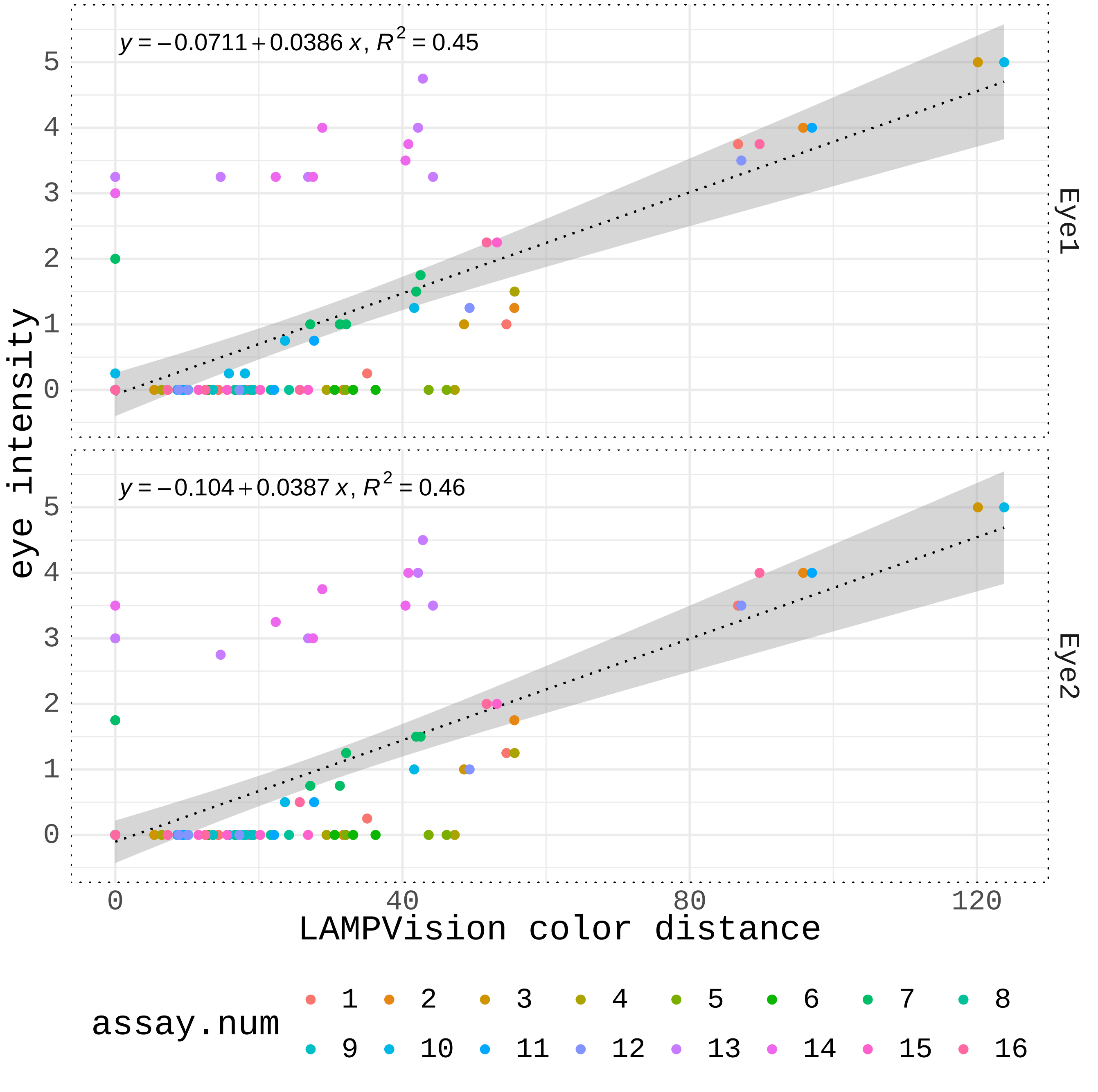

### Fig3c.png

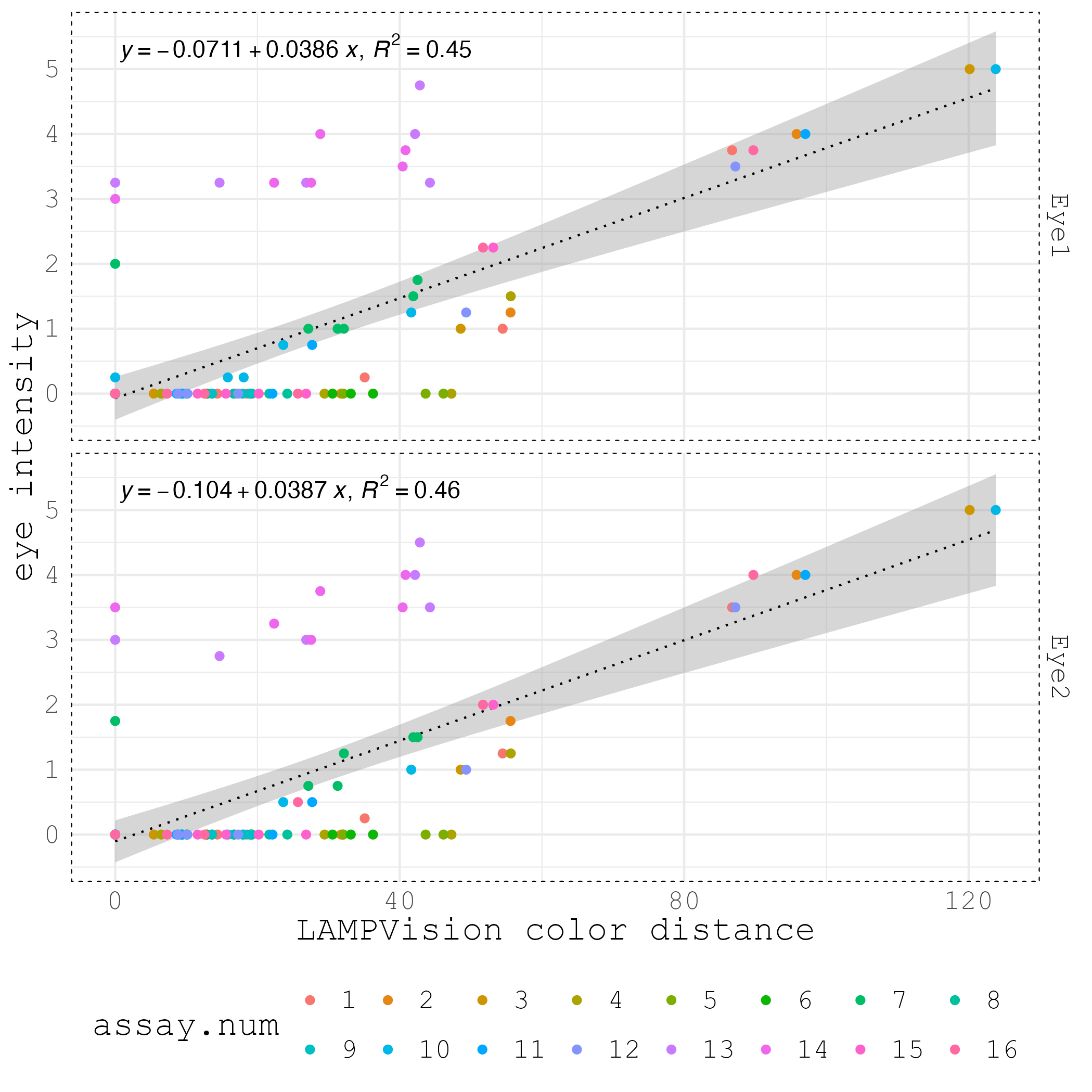

### Fig3d.pdf

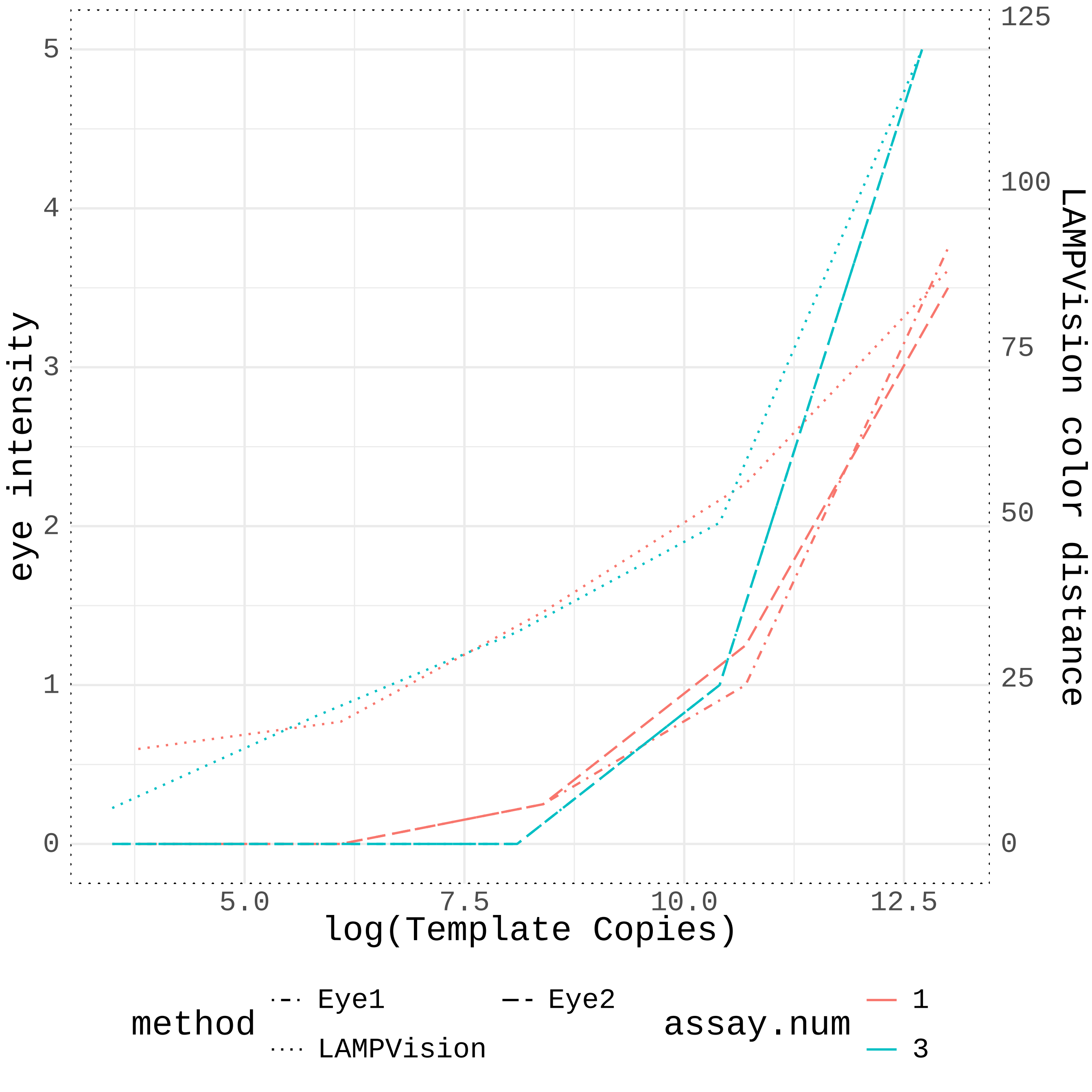

### Fig3d.png

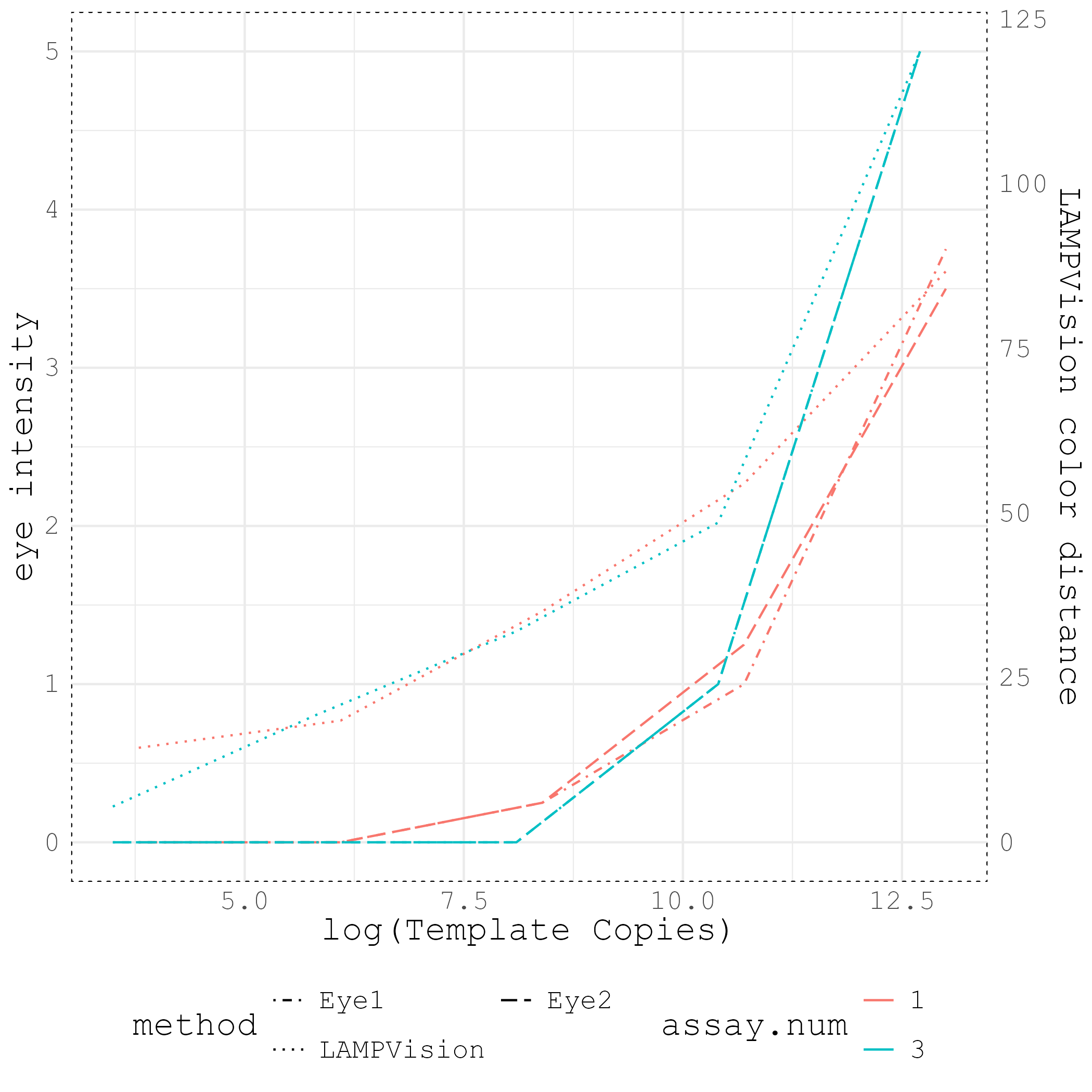

### Rep1.circles.jpg

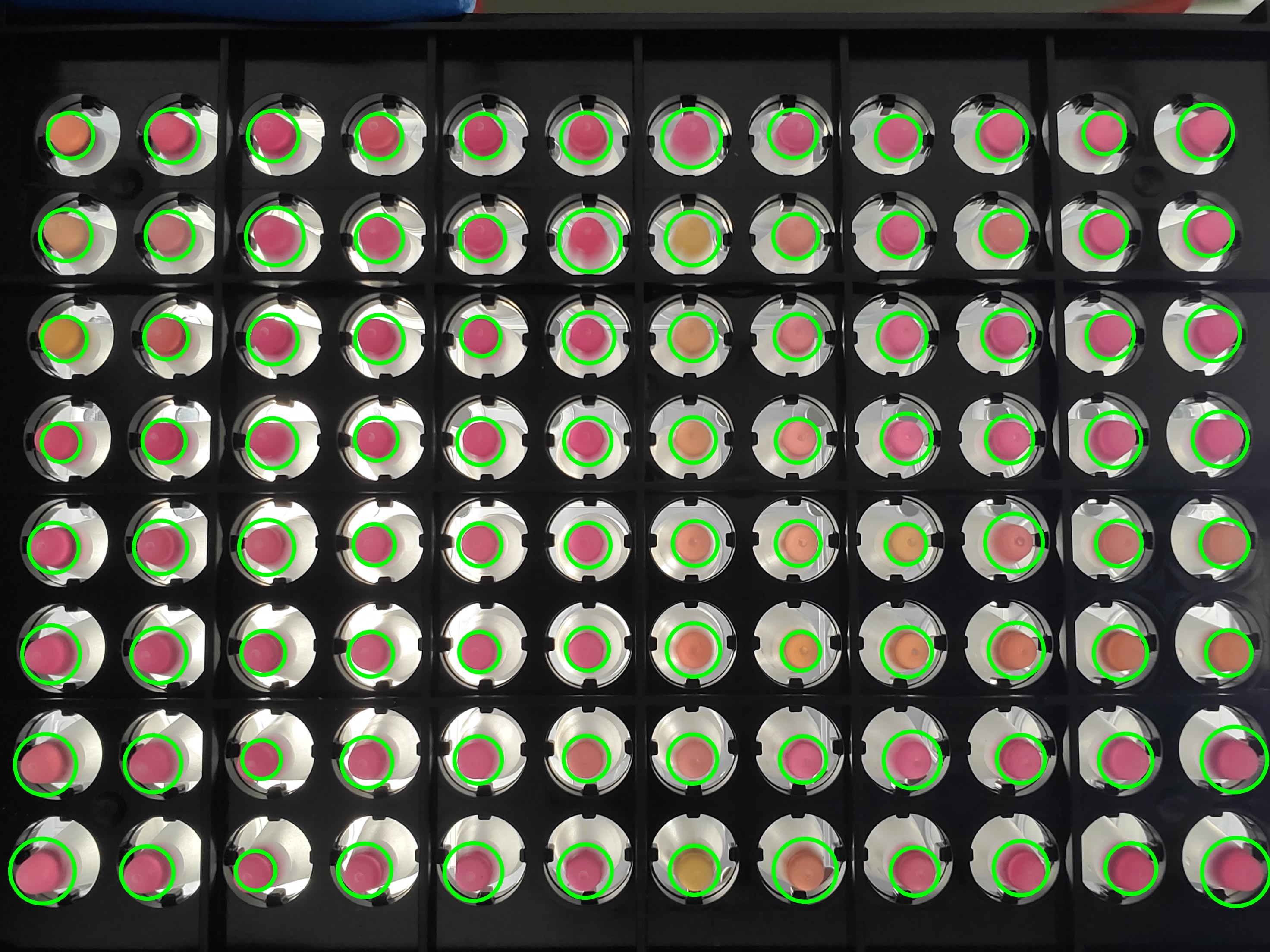

### Rep1.jpg

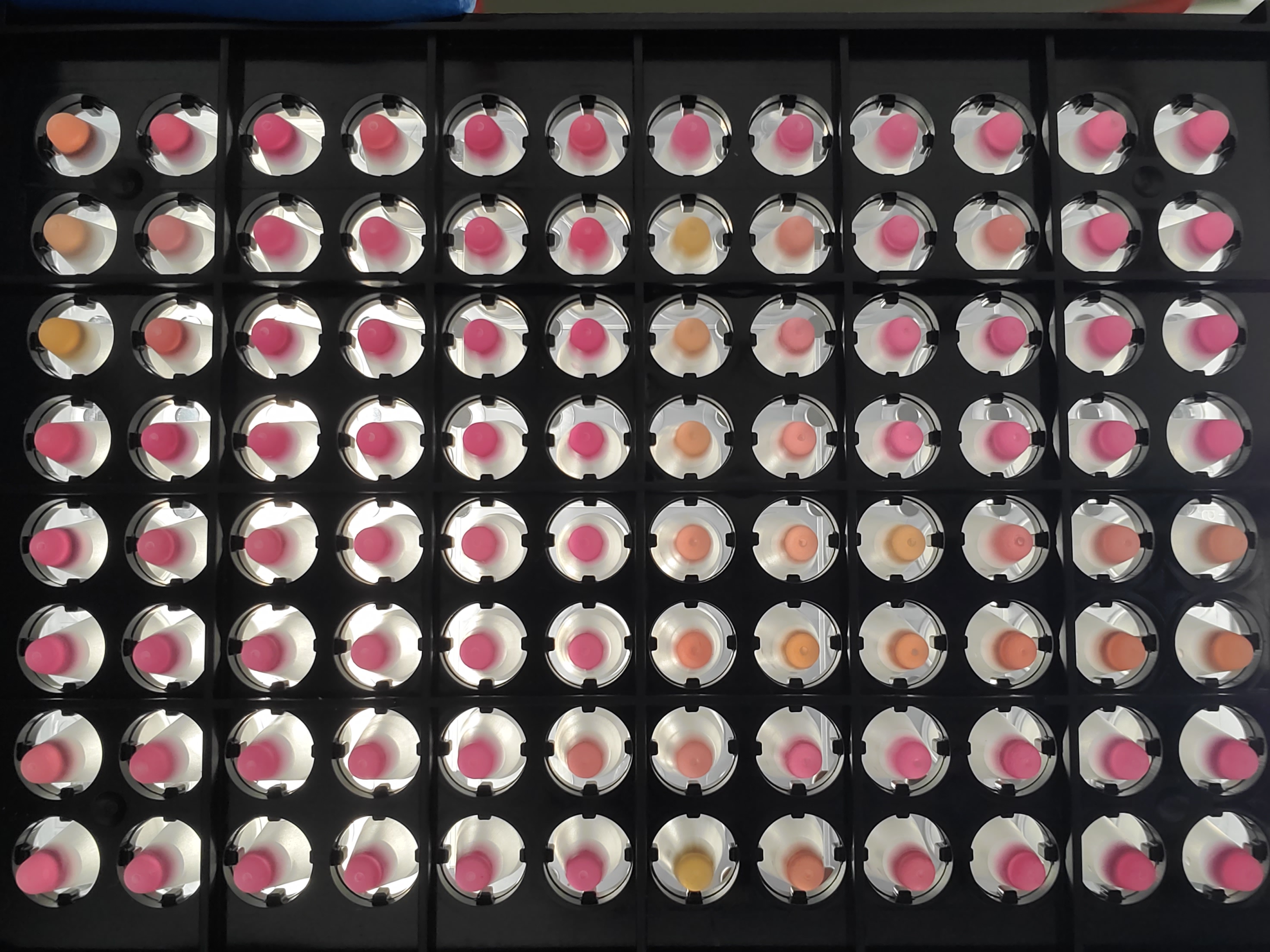

### Rep2.circles.jpg

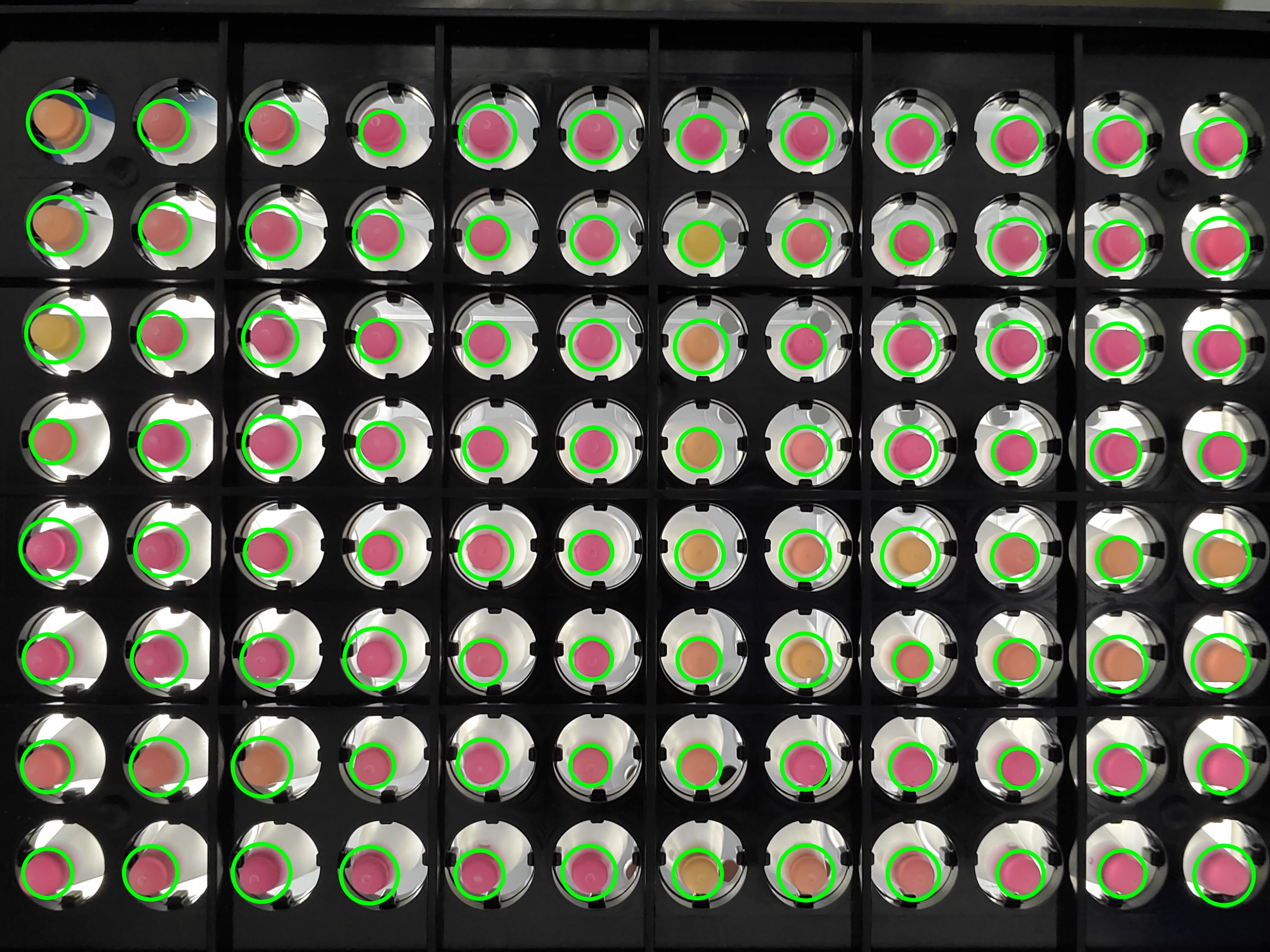

### Rep2.jpg

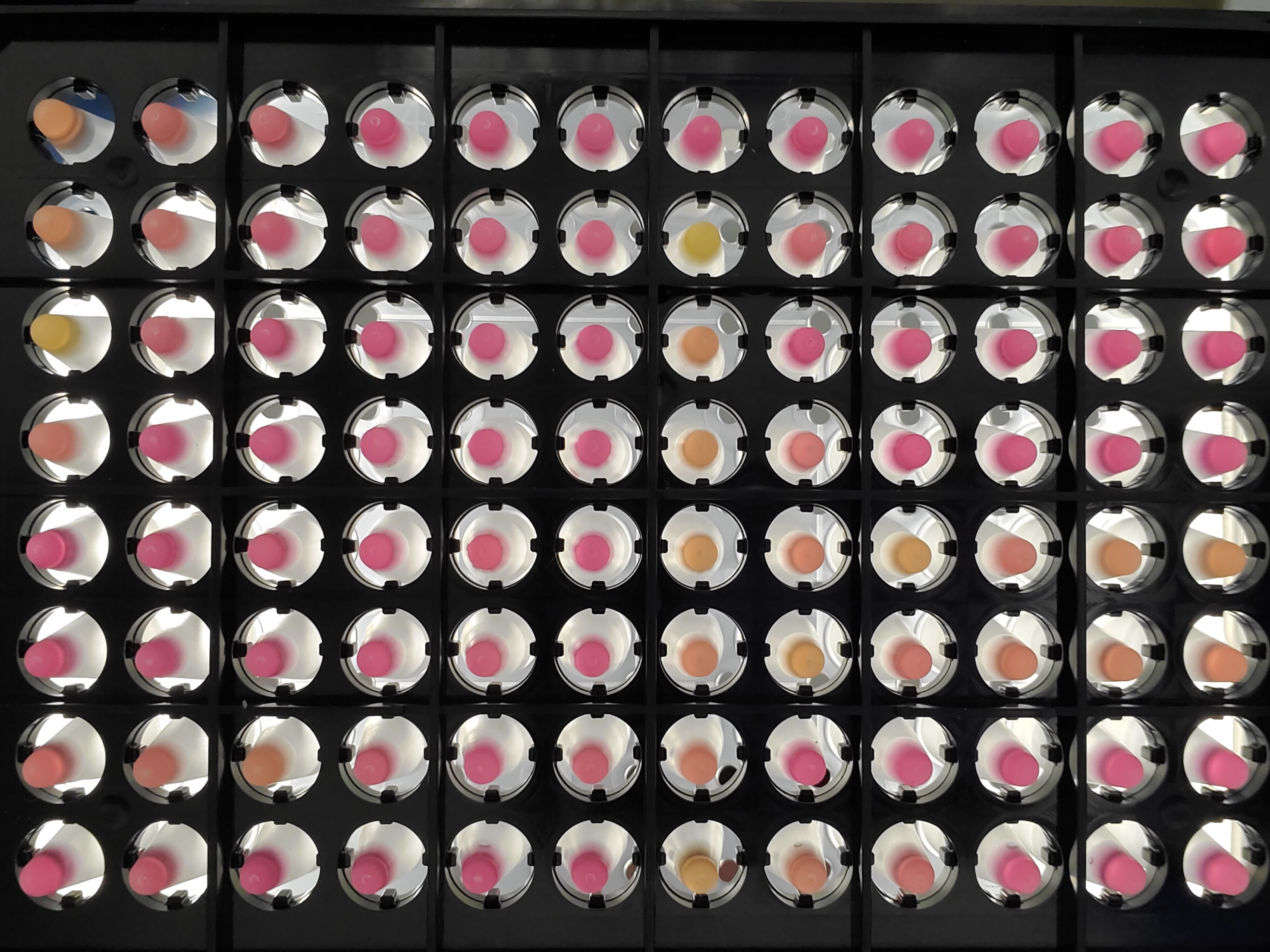

### Rep3.circles.jpg

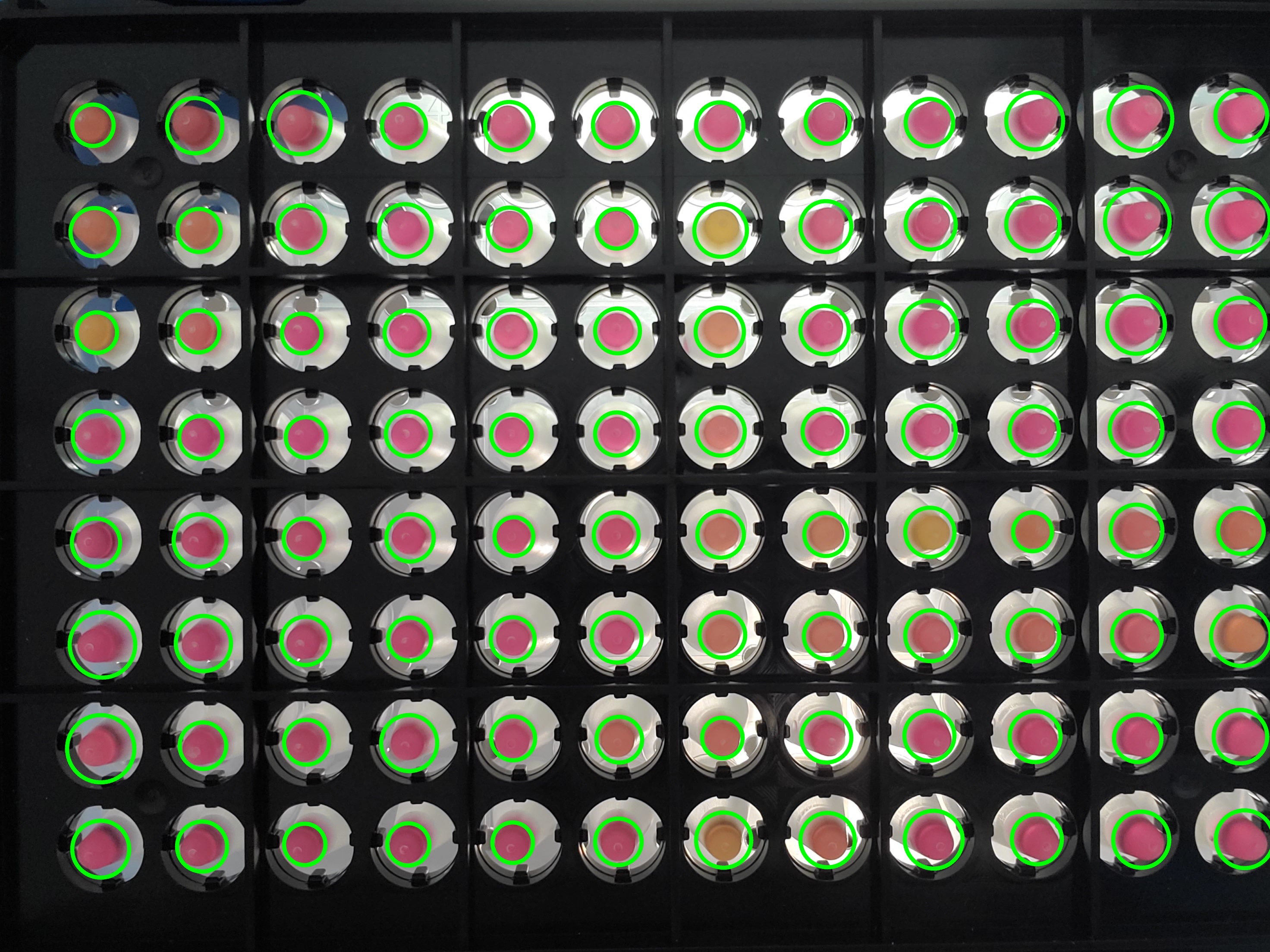

### Rep3.jpg

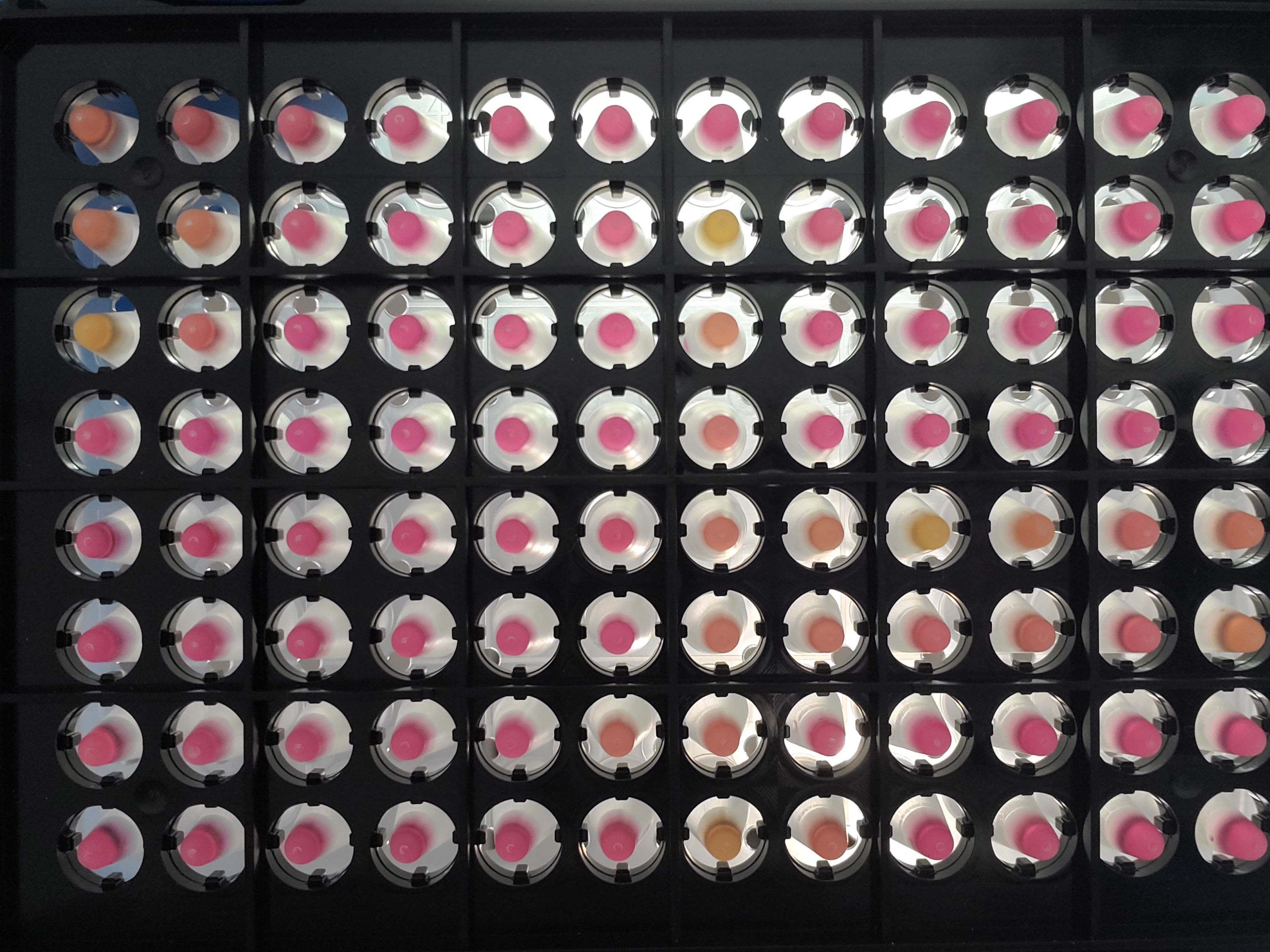

### Rep4.circles.jpg

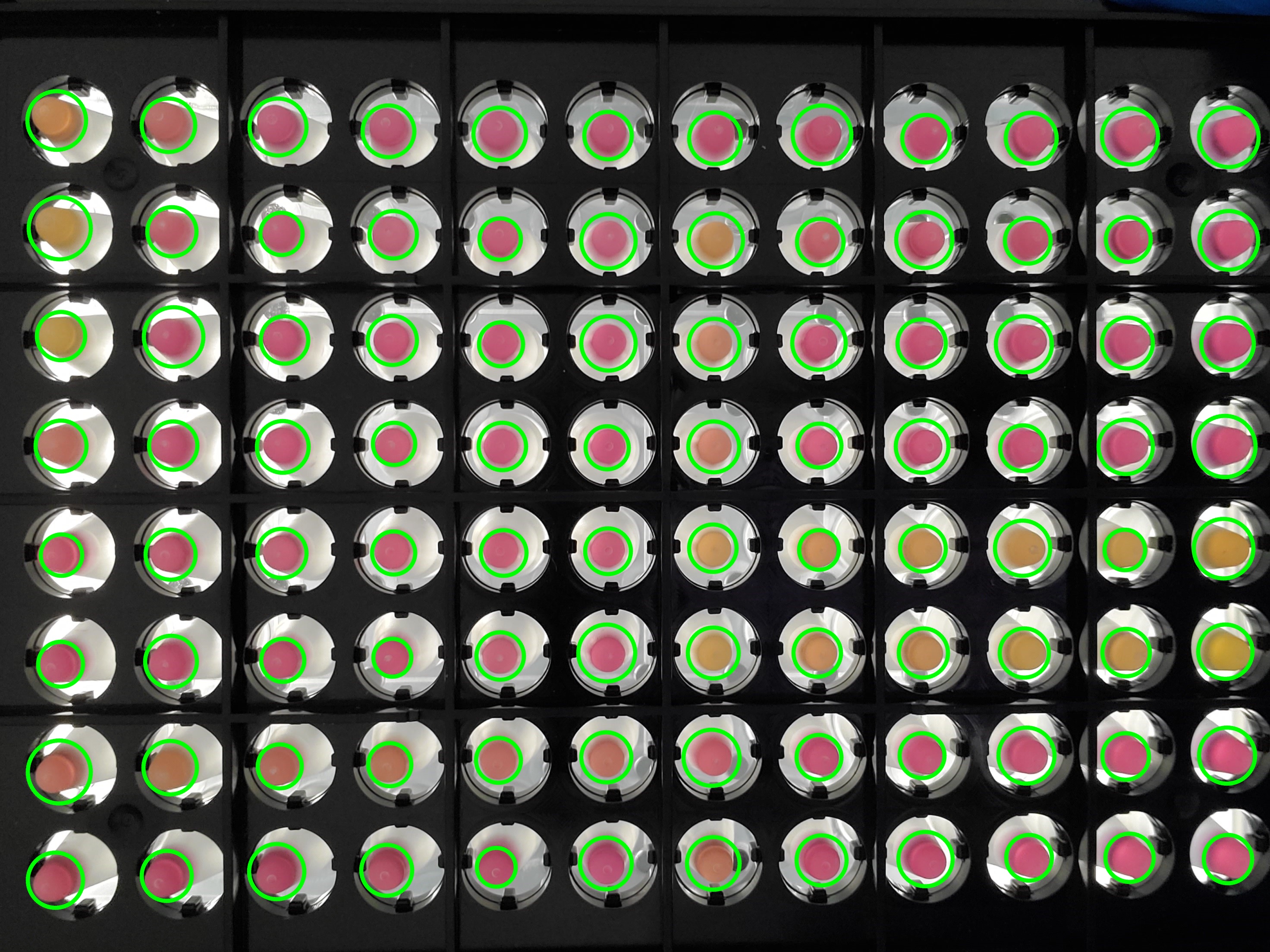

### Rep4.jpg

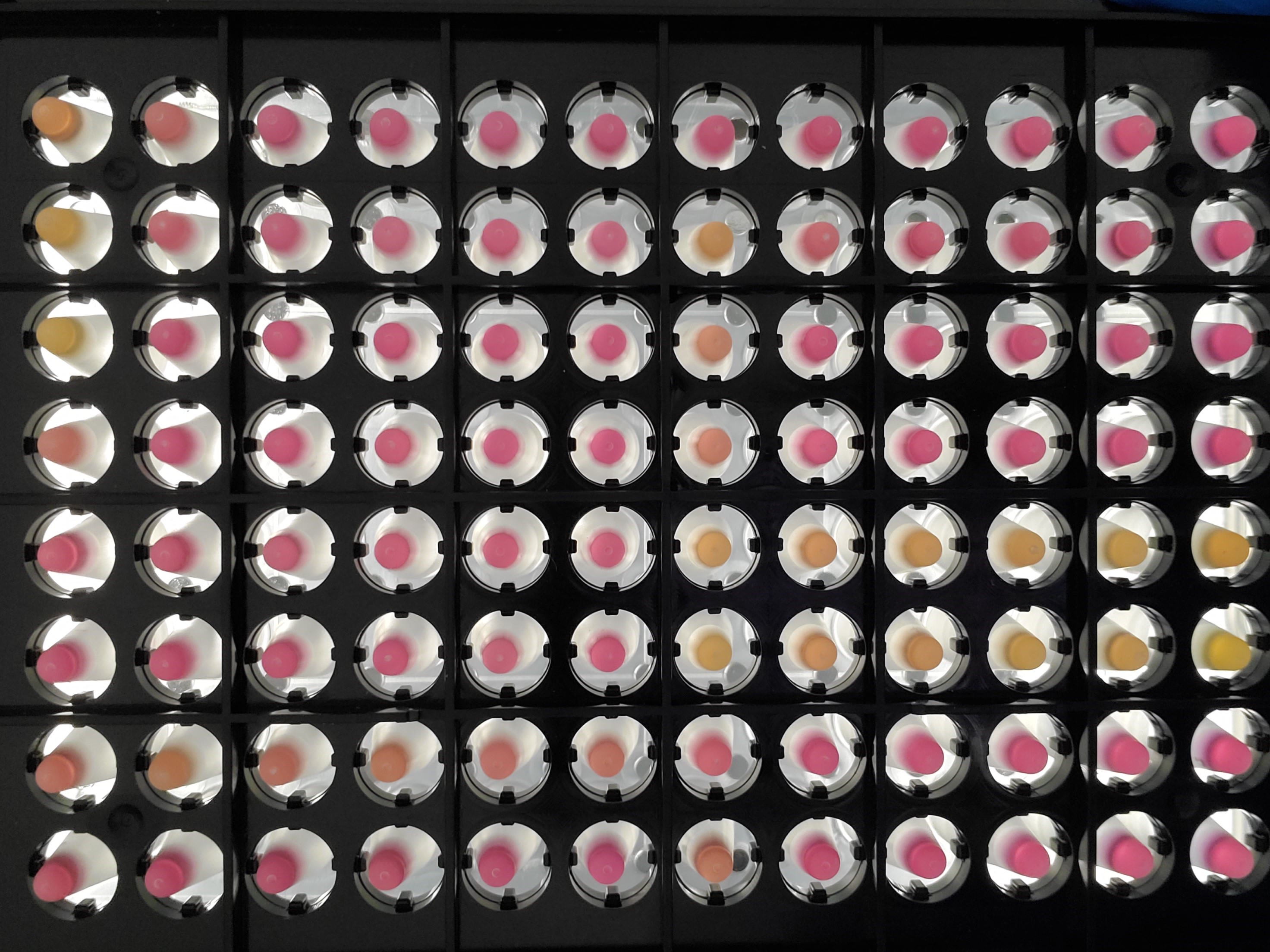

### Rplots.pdf

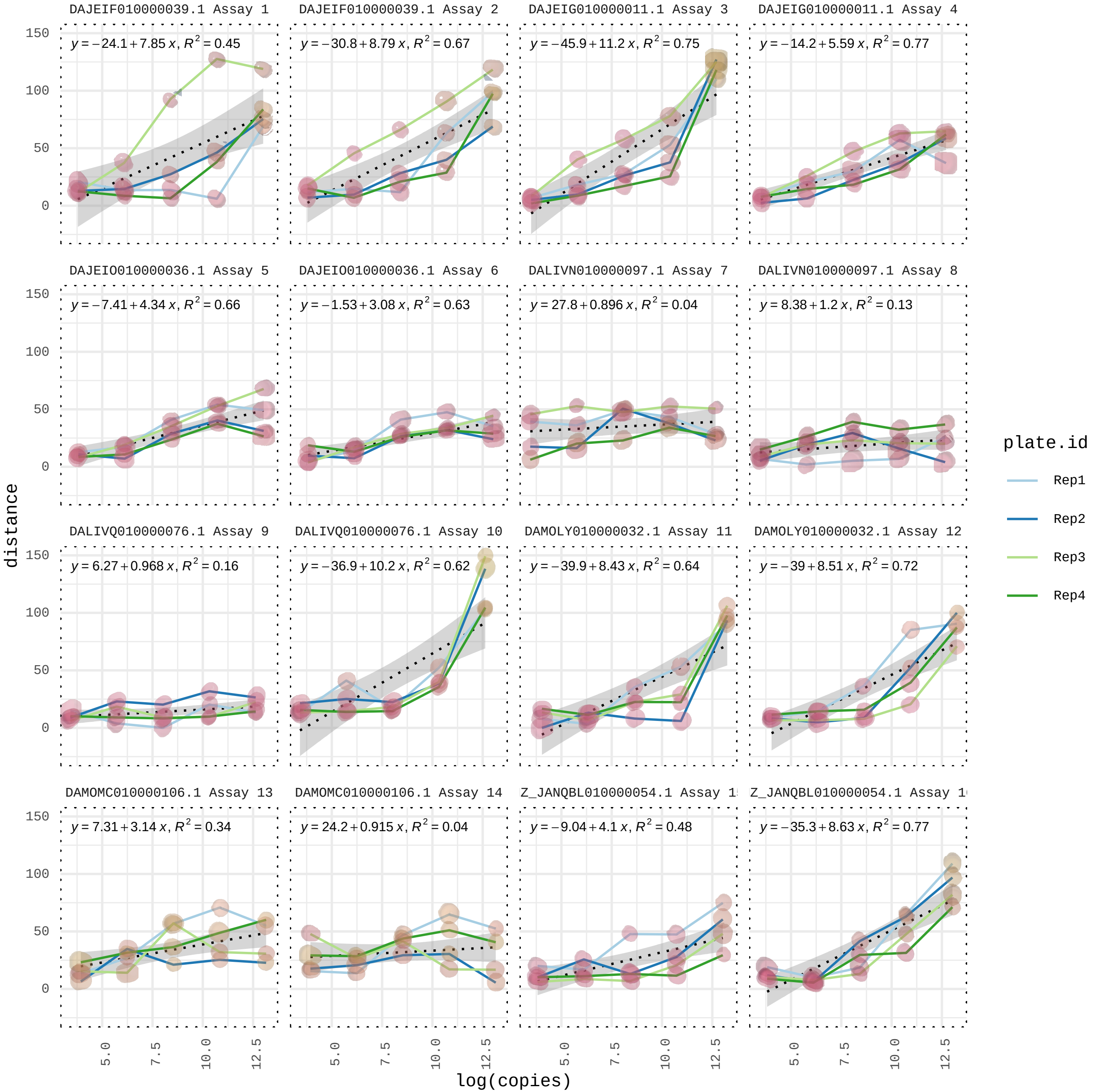

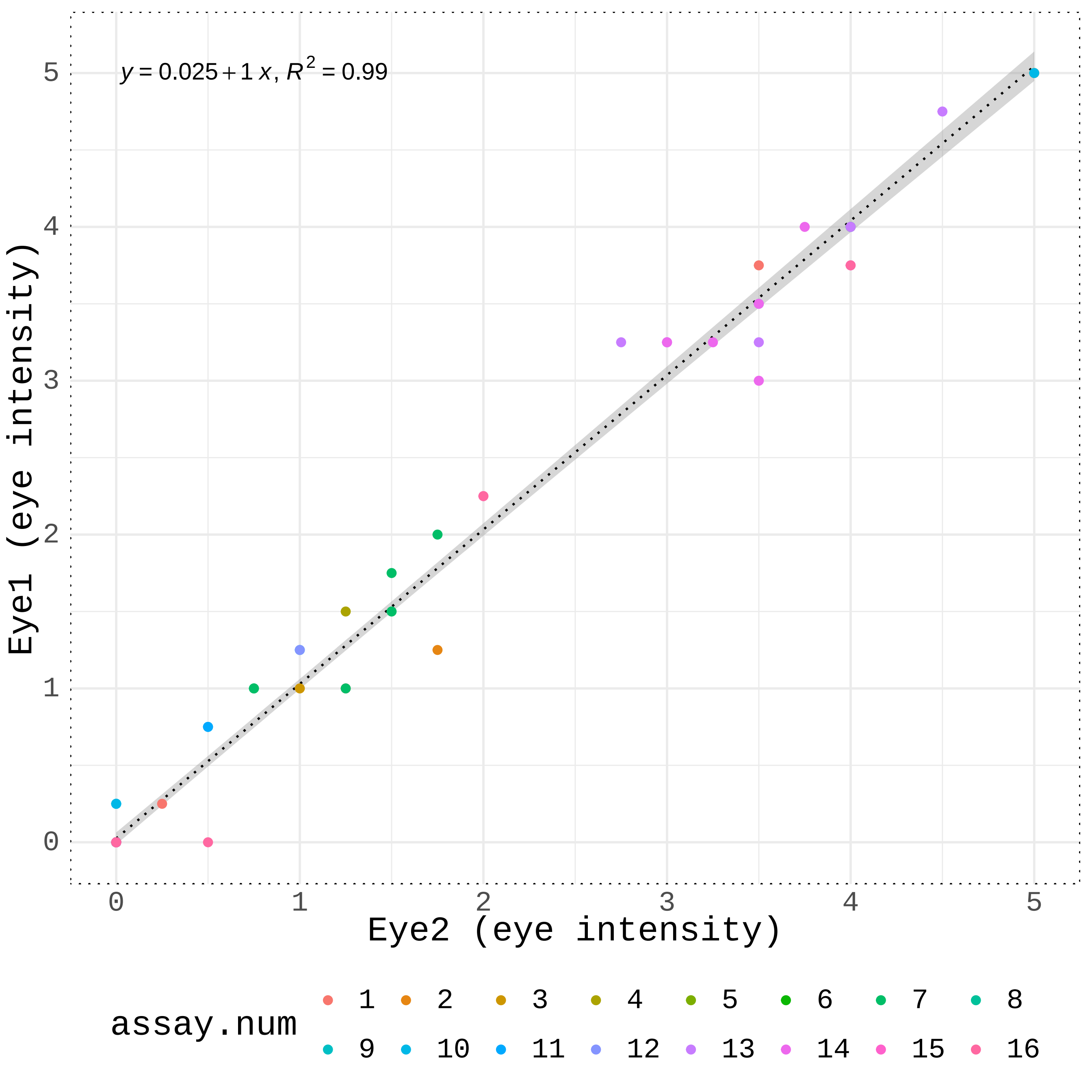

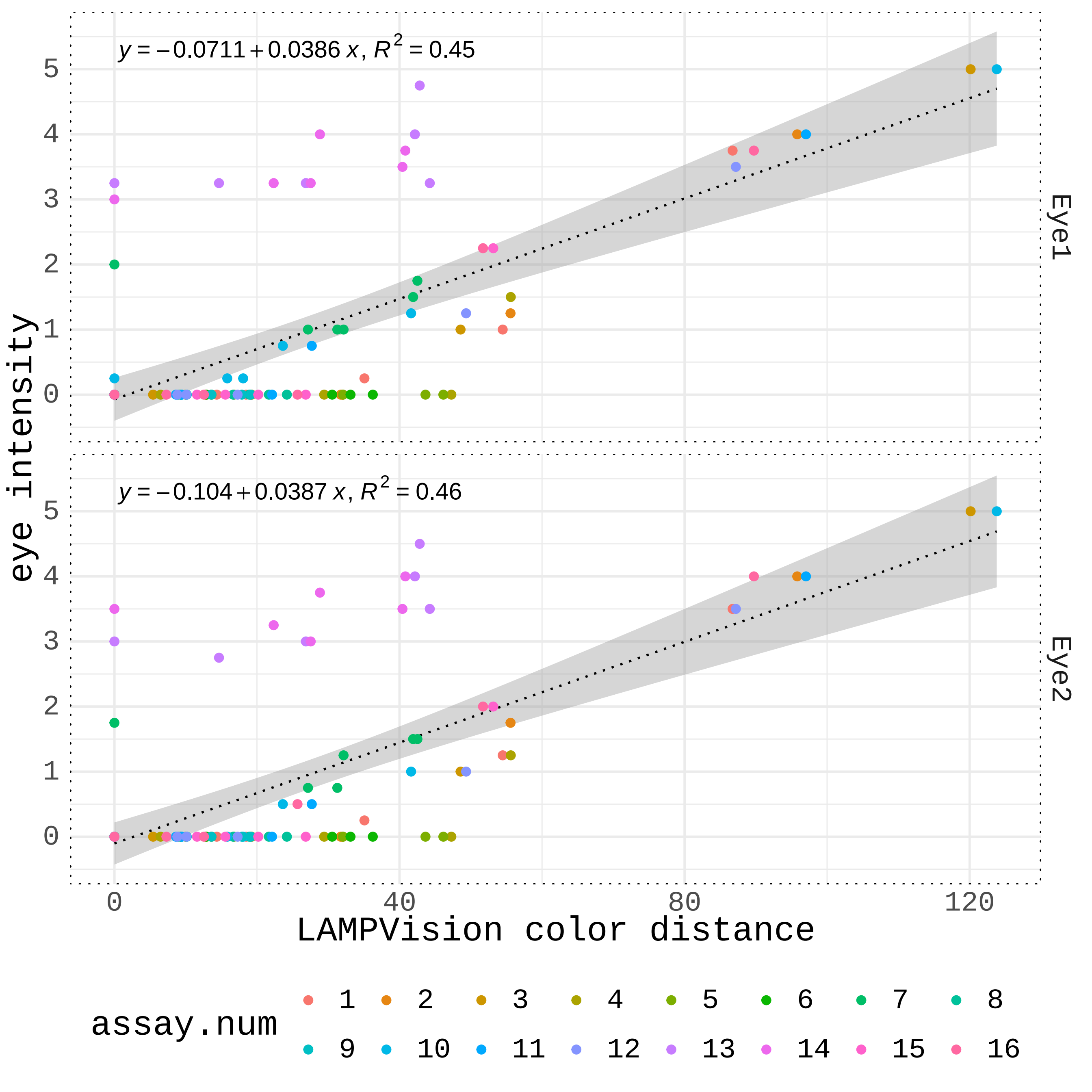

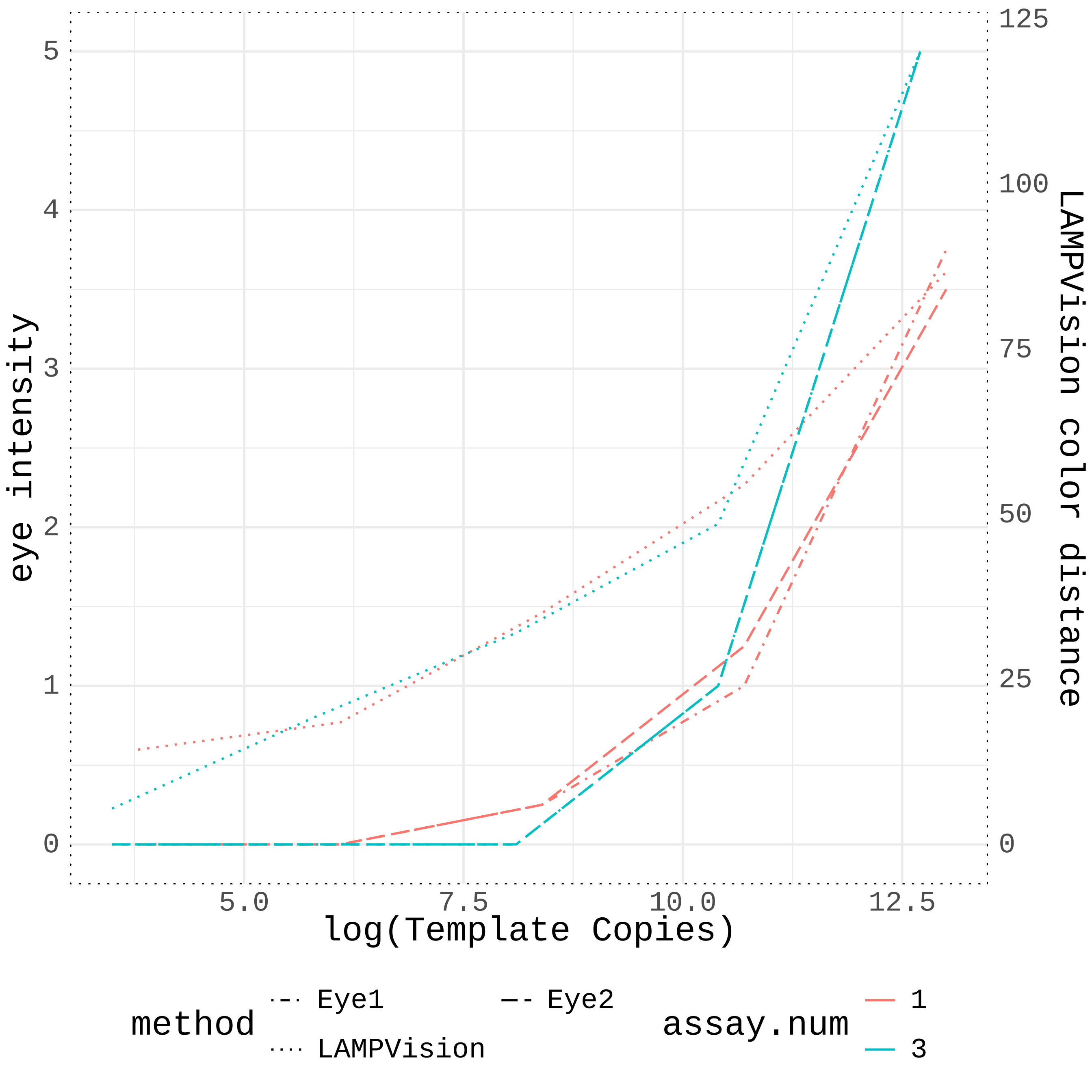
